# Axon guidance pathways modulate neurotoxicity of ALS-associated UBQLN2

**DOI:** 10.1101/2022.10.31.514355

**Authors:** Sang Hwa Kim, Kye D. Nichols, Eric N. Anderson, Yining Liu, Nandini Ramesh, Weiyan Jia, Connor J. Kuerbis, Mark Scalf, Lloyd M. Smith, Udai Bhan Pandey, Randal S. Tibbetts

**Author notes:** Corresponding authors: Sang Hwa Kim, Randal Tibbetts.

## Abstract

Mutations in the ubiquitin (Ub) chaperone *Ubiquilin 2 (UBQLN2)* cause X-linked forms of amyotrophic lateral sclerosis (ALS) and frontotemporal dementia (FTD) through unknown mechanisms. Here we show that aggregation-prone, ALS-associated mutants of UBQLN2 (UBQLN2^ALS^) trigger heat stress-dependent neurodegeneration in Drosophila. A genetic modifier screen implicated endolysosomal and axon guidance genes, including the netrin receptor, Unc-5, as key modulators of UBQLN2 toxicity. Reduced gene dosage of *Unc-5* or its coreceptor *Dcc/frazzled* diminished neurodegenerative phenotypes, including motor dysfunction, neuromuscular junction defects, and shortened lifespan, in flies expressing UBQLN2^ALS^ alleles. Induced pluripotent stem cells (iPSCs) harboring UBQLN2^ALS^ knockin mutations exhibited lysosomal defects while inducible motor neurons (iMNs) expressing UBQLN2^ALS^ alleles exhibited cytosolic UBQLN2 inclusions, reduced neurite complexity, and growth cone defects that were partially reversed by silencing of UNC5B and DCC. The combined findings suggest that altered growth cone dynamics are a conserved pathomechanism in UBQLN2-associated ALS/FTD.

## Introduction

### Ubiquilins and proteostasis

Defective protein folding and proteostatic stress are common pathogenic mechanisms linking genetically and anatomically diverse neurodegenerative diseases^1^. The steady state levels—and ultimately neurotoxicity—of aggregation-prone proteins are determined through a balance of protein production and protein clearance^2^. The convergence of aging-dependent declines in protein degradation and nuclear import with environmental stresses may push this equation toward irreversible protein aggregation that sets the stage for neurodegenerative processes^3^. Thus, enhancing protein degradation capacity—or reducing the aggregation potential of aggregation-prone proteins—represents a promising therapeutic avenue for ALS and other neurodegenerative proteinopathies.

The highly conserved ubiquilin (UBQLN) gene family fulfills diverse roles in protein folding, shuttling, and degradation^4,5^. All eukaryotic UBQLNs feature an amino-terminal ubiquitin-like (UBL) domain and a carboxyl-terminal ubiquitin-associated separated by a low complexity, methionine-rich central region harboring variable numbers of STI1-like repeats first identified in the yeast stress-inducible 1 (Sti1) protein and later described in several distinct classes of protein chaperones ^6-10^. Mammals encode four ubiquilin proteins of which three (UBQLN1, UBQLN2, and UBQLN4) are widely expressed. UBQLN1 and UBQLN2 share >70% amino acid identity and are presumed to function in semi-redundant fashion, whereas UBQLN4 is a more distantly related paralog.

Although specific functions of individual ubiquilins in mammals are largely unknown, a generic model for UBQLN function holds that the UBA domain engages ubiquitylated substrate while the UBL domain engages the proteasome, leading to substrate degradation^11^. STI1-like repeats form a hydrophobic groove that is thought to engage hydrophobic regions of client proteins^6^. It has been reported that UBQLN deficiency leads to defects in autophagy and ER-associated protein degradation (ERAD)^12-15^. The central methionine-rich domain has been shown to bind transmembrane domains of mitochondrial proteins, which appears central to their UBQLN-dependent triage and degradation^11^.

### UBQLN2 mutations in ALS/dementia

Interest in ubiquilin function was greatly stimulated by the discovery that dominant mutations in UBQLN2 cause X-linked ALS/frontotemporal dementia (FTD)^16,17^. Most ALS-associated mutations in UBQLN2 are clustered within 42-amino acid proline-rich repeat (PRR) that is unique to UBQLN2^16^; however, disease-linked mutations outside this region have also been described^18,19^. In addition, UBQLN2-mutant patients exhibit a range of phenotypes that includes FTD, ALS, and spastic paraplegia^20^. Interestingly, ubiquilin-positive aggregates are a near universal occurrence in TDP-43-positive ALS, as well as ALS linked to *C9ORF72* expansions (C9-ALS)^21^. These correlative findings suggest that ubiquilin pathology may contribute to the molecular pathogenesis of ALS even in the absence of *UBQLN2* gene mutations.

Transgenic or virus-directed expression of UBQLN2^ALS^ mutants in rodents recapitulates protein UBQLN2 inclusions seen in ALS/FTD patients and elicits variable phenotypic abnormalities ranging to mild gait and memory defects to neuronal loss, paralysis, and early death^22-26^. Coexpression of UBQLN2^P497H^ and an ALS-associated TDP-43 allele under control of the neurofilament heavy gene promoter caused severe motor neuron loss and muscle wasting^27^. Brain-directed expression of wild-type UBQLN2 also caused toxicity phenotypes^24-26^ potentially due to disruptions in Ub homeostasis. Finally, like many proteins implicated in ALS/FTD^28^, UBQLN2 harbors low complexity regions and undergoes liquid-liquid phase separation^29,30^. While ALS-associated mutations in the PRR may interfere with UBQLN2 liquid demixing, the relevance to disease pathogenesis is presently unclear.

In previous work we reported that expression of ALS-associated *UBQLN2* mutants caused mutation-dependent neurotoxicity in Drosophila^31^. Here, we carried out deficiency screens for UBQLN2 toxicity modifiers using flies expressing UBQLN2^ALS^ alleles with differing aggregation potential. Suppressor genes emerging from this screen were then tested for impacts on the toxicity of endogenous UBQLN2^ALS^ mutants in iMNs. Our findings suggest that endolysosomal dysfunction and axon guidance defects are phenotypic drivers of neurodegeneration in UBQLN2-associated ALS/FTD.

## Results

### An aggregation-prone UBQLN2^4XALS^ allele exhibits heat stress (HS)-dependent eye toxicity

We previously reported that two copy expression of clinical UBQLN2^ALS^ alleles caused mild eye toxicity when expressed in the Drosophila compound eye at 23°C under the control of eye-specific GMR driver (GMR>UBQLN2 flies)^31^. Reasoning that UBQLN2-associated phenotypes may be worsened by heat stress (HS) we compared eye morphology in GMR>UBQLN2 flies reared at 22°C and 29°C during all developmental stages. While none of the single-copy UBQLN2 transgenes caused an overt external eye phenotype at 22°C, flies expressing a combinatorial, aggregation-prone, UBQLN2^4XALS^ mutant at 29°C (Figure 1A), exhibited a moderately severe rough eye (RE) phenotype that was characterized by eye depigmentation and loss of ommatidial facets in both male and female flies (Figure 1B).

**Figure 1.**
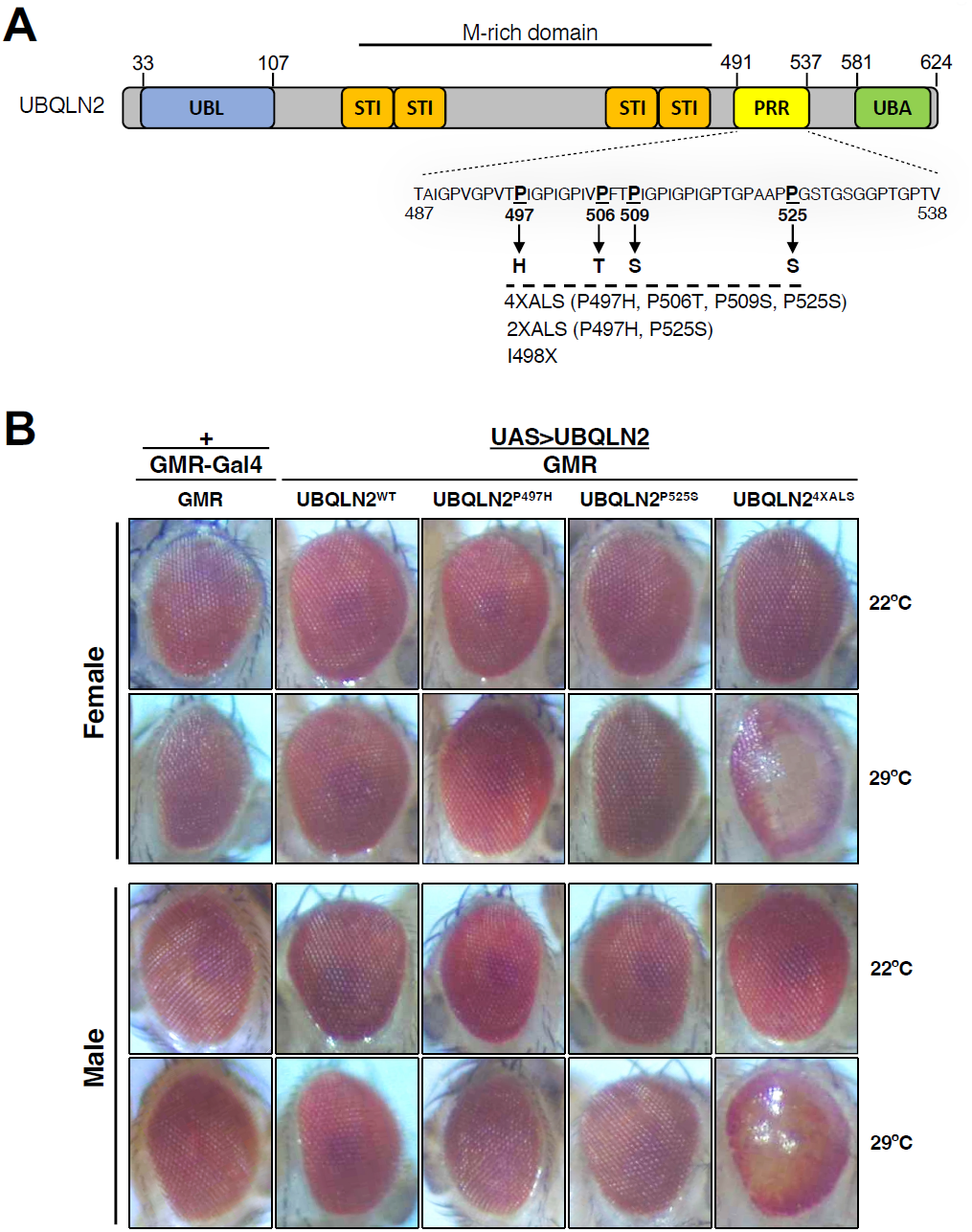
UBQLN2 exerts heat-stress-dependent toxicity. **(A)** Schematic of UBQLN2 and ALS-associated mutations. Approximate locations of Ub-like (UBL); STI1-like (STI); proline rich repeat (PRR); and Ub-associated (UBA) domains are shown, as are ALS-associated mutations investigated in this study. (**B)** Eye images from flies expressing UBQLN2^WT^, UBQLN2^P497H^, UBQLN2^P525S^ or UBQLN2^4XALS^ under control of the eye-specific GMR driver at 22°C and 29°C. Note depigmentation and destruction of ommatidial facets in UBQLN2^4XALS^ flies reared at 29°C.

To determine whether HS affected UBQLN2 aggregation, we measured detergent solubility of UBQLN2^WT^ and UBQLN2^ALS^ mutants expressed under control of GMR. This revealed that the protein insolubility of UBQLN2^4XALS^ in whole fly heads was not further exacerbated by HS (Figure supplement 1A). On the other hand, an F594A mutation that abolished Ub binding of UBQLN2^31^ dramatically decreased intraneuronal aggregation of UBQLN2^4XALS^ and reversed UBQLN2^4XALS^-mediated eye degeneration at 29°C (Figure supplement 1B, C). We also performed quantitative mass spectrometry (MS) to assess relative abundance of UBQLN2^WT^, UBQLN2^4XALS^, and endogenous Drosophila Ubqln (dUbqln) in whole-head extracts. The average number of peptide spectral matches (PSMs) for hUBQLN2 were comparable between GMR>UBQLN2^WT^ and GMR>UBQLN2^4XALS^ flies reared at 22°C and 29°C. As expected, absolute numbers of hUBQLN2 peptides were higher in GMR>UBQLN2^WT^ and GMR>UBQLN2^4XALS^ flies reared at 29°C owing to heat-inducibility of the UAS regulatory region. hUBQLN2-unique peptides were ∼6-fold more abundant than dUbqln peptides in both GMR>UBQLN2^WT^ and GMR>UBQLN2^4XALS^ heads, providing a lower limit of hUBQLN2 overexpression (Figure supplement 1D). These findings suggest that enhanced phenotypes of GMR>UBQLN2^4XALS^ flies relative to GMR>UBQLN2^WT^ flies are not solely due to differences in protein expression.

### Transcriptomic analysis of UBQLN2^ALS^ flies

We next performed RNA-Seq analysis of whole heads from GMR>UBQLN2^WT^, GMR>UBQLN2^P497H^ and GMR>UBQLN2^4XALS^ flies. Consistent with their severe eye phenotype, GMR>UBQLN2^4XALS^ flies exhibited a distinct gene expression signature relative to GMR>UBQLN2^WT^ and GMR>UBQLN2^P497H^ flies, which clustered together in principle component analysis (PCA) (Figure 2A). Using an FDR of <.05, 629 genes showed selective differential expression between GMR>UBQLN2^4XALS^ and GMR>UBQLN2^WT^ flies at 29°C (Figure supplement 2B, C). Of these, 402 genes were upregulated and 227 genes were downregulated in GMR>UBQLN2^4XALS^ flies (Supplementary Datasets 1 and 2, respectively). Gene ontology (GO) analysis revealed that genes involved in eye development and phototransduction, including ninaA, ninaE, and Rh3, were broadly downregulated in GMR>UBQLN2^4XALS^ flies, likely reflecting degenerative cell loss (Figure supplement 2D and Supplementary Dataset 2). Upregulated GO terms included response to biotic stimulus, which includes Drosophila innate immunity genes such as Dro, AttB, AttC and response to UV light, which includes the small heat-shock proteins HSP23, HSP26, and HSP27. (Figure 2D and Supplementary Dataset 1). Upregulation of HSPs may be driven by UBQLN2^4XALS^ misfolding. In addition, 602 genes were differentially expressed between GMR>Gal4 and GMR>UBQLN2^WT^ flies indicating that UBQLN2 overexpression has a substantial impact on gene expression, while only 64 genes were differentially expressed between GMR>UBQLN2^WT^ and GMR>UBQLN2^P497H^ flies. The highly overlapping gene expression profiles between GMR>UBQLN2^WT^ and GMR>UBQLN2^P497H^ flies is consistent with their qualitatively similar eye phenotypes. Finally, 28 differentially expressed genes were common to GMR>UBQLN2^4XALS^ and GMR>UBQLN2^P497H^ flies relative to GMR>UBQLN2^WT^ flies (Figure supplement 2E). Overall, these findings reveal that the strong eye phenotype of GMR>UBQLN2^4XALS^ is accompanied by robust changes in gene expression.

**Figure 2.**
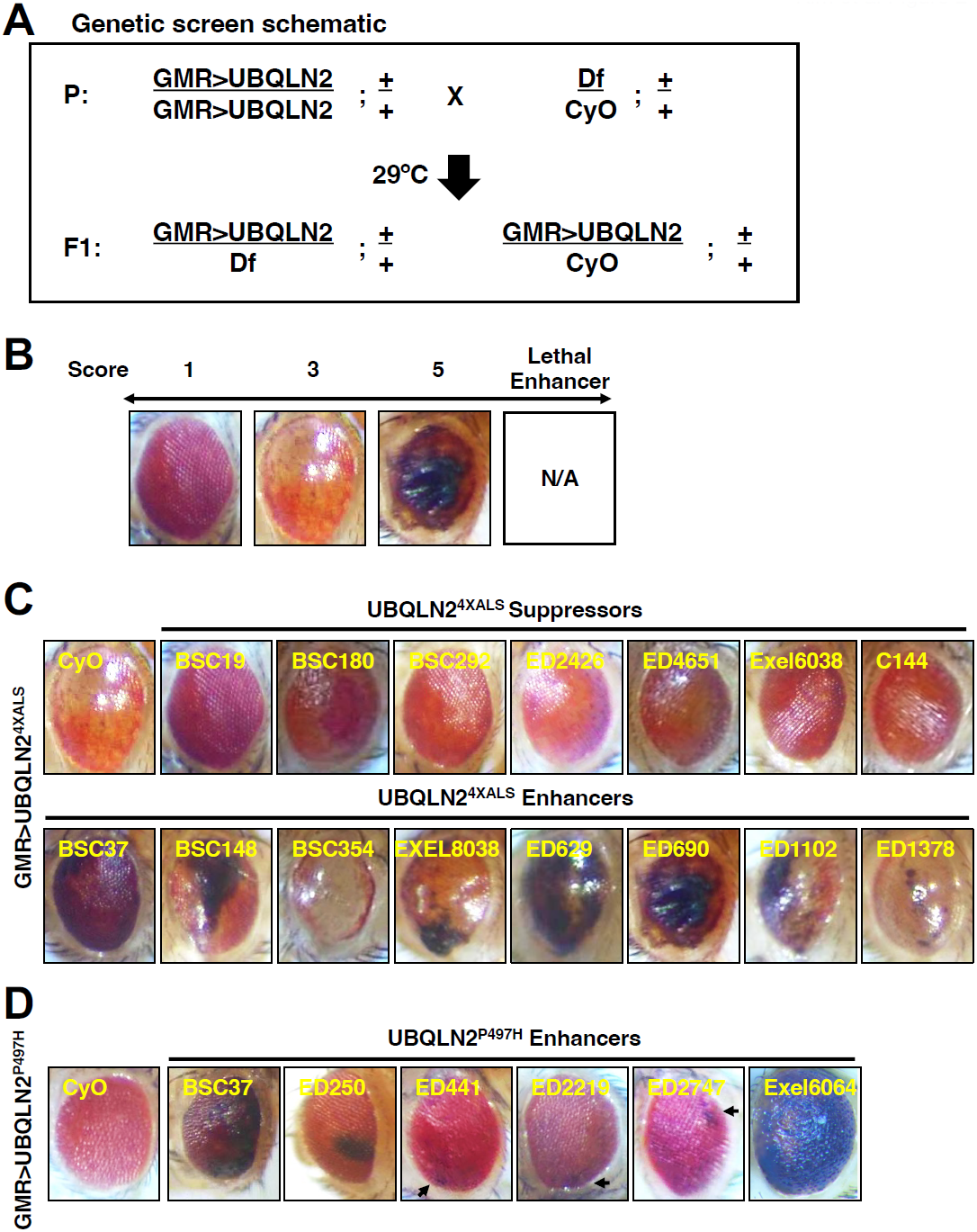
Identification of UBQLN2 modifier genes. **(A)** Schematic of deficiency (Df) screen for UBQLN2 modifiers. **(B)** Representative eye images and scoring rubric for F1 progeny of GMR>UBQLN2^4XALS^ flies crossed to Df lines. **(C)** Representative UBQLN2^4XALS^ modifier genes. Eye images were taken of F1 progeny from crosses of GMR>UBQLN2^4XALS^ to indicated Df lines at 1-3 day post eclosion. Suppressors and enhancers are shown in top and bottom rows, respectively. **(D)** Representative eye images of UBQLN2^P497H^ enhancers. Arrows indicate foci of eye degeneration.

### Deficiency screens for UBQLN2^ALS^ modifier genes

The RE phenotype of GMR>UBQLN2^4XALS^ flies allowed us to perform genetic modifier screens. To this end, we screened a Bloomington deficiency (Df) library of 194 lines spanning 85% of chromosome 2 (Figure 2A). The screen was carried out at 29°C and eye phenotypes were scored on a scale of 1-5, with a score of “1” representing a morphologically normal eye; a score of “3” representing the unmodified UBQLN2^4XALS^ eye phenotype; and a score of “5” representing eyes harboring >50% necrotic patches (Figure 2B). Lethal enhancers were also noted. While the mild eye phenotype of GMR>UBQLN2^P497H^ (score 1) flies largely precluded identification of suppressors, the side-by-side screening of this line allowed us to identify shared and/or mutation-specific enhancers. Candidate modifier Dfs were subjected to secondary screens against UBQLN2^WT^, UBQLN2^P497H^, and UBQLN2^4XALS^. The UBQLN2^4XALS^ screen identified seven suppressors, three of which overlapped a common genomic interval, and 23 enhancers, including 15 lethal enhancers (Figure 2C-Table supplement 1). The UBQLN2^P497H^ screen identified 6 enhancers, two of which were also identified in the UBQLN2^4XALS^ screen (Figure 2D-Table supplement 1). All Dfs that enhanced the GMR>UBQLN2^P497H^ eye phenotype also caused enhanced eye phenotypes in GMR>UBQLN2^WT^ flies.

A combination of iterative Df screening and RNAi screening was then used to map causal genes within three UBQLN2^4XALS^ suppressor loci (BSC180, ED2426 and Exel6038) and one enhancer locus common to UBQLN2^4XALS^ and UBQLN2^P497H^ (BSC37) (Table supplement 1). Among 17 annotated genes within BSC37 (Figure 3A), we prioritized *Rab5*, which encodes an early endosomal protein that interacts with the ALS2 gene product, Alsin^32,33^. In support of a role for Rab5 as a UBQLN2 modifier gene, Rab5 knockdown phenocopied the hyperpigmented phenotype seen in GMR>UBQLN2^4XALS^ flies crossed to BSC37 at 29°C (Figure 3B, C). Rab5 knockdown also caused a hyperpigmented eye phenotype in GMR>UBQLN2^WT^ and GMR>UBQLN2^P497H^ flies (Figure 3C), indicating Rab5 is a mutation-independent UBQLN2 enhancer. Interestingly, while Rab5 knockdown also caused hyperpigmented eye patches in GMR>UBQLN2^WT^, and GMR>UBQLN2^P497H^ flies reared at 22°C, GMR>UBQLN2^4XALS^/shRab5 flies reared at 22°C failed to exhibit eye patches (Figure 3C). This finding is consistent with the idea that UBQLN2^4XALS^ phenotypes are HS dependent. Finally, we showed that overexpression of GFP-Rab5 partially rescued the eye phenotype of GMR>UBQLN2^WT^, GMR>UBQLN2^P497H^, and GMR>UBQLN2^4XALS^ flies reared at 29°C (Figure 3D). Rab5 knockdown or overexpression did not affect UBQLN2 expression (Figure 3E). The combined findings suggest that Rab5 is a general modulator of UBQLN2 toxicity and that UBQLN2 overexpression interferes with endolysosomal function in flies.

**Figure 3.**
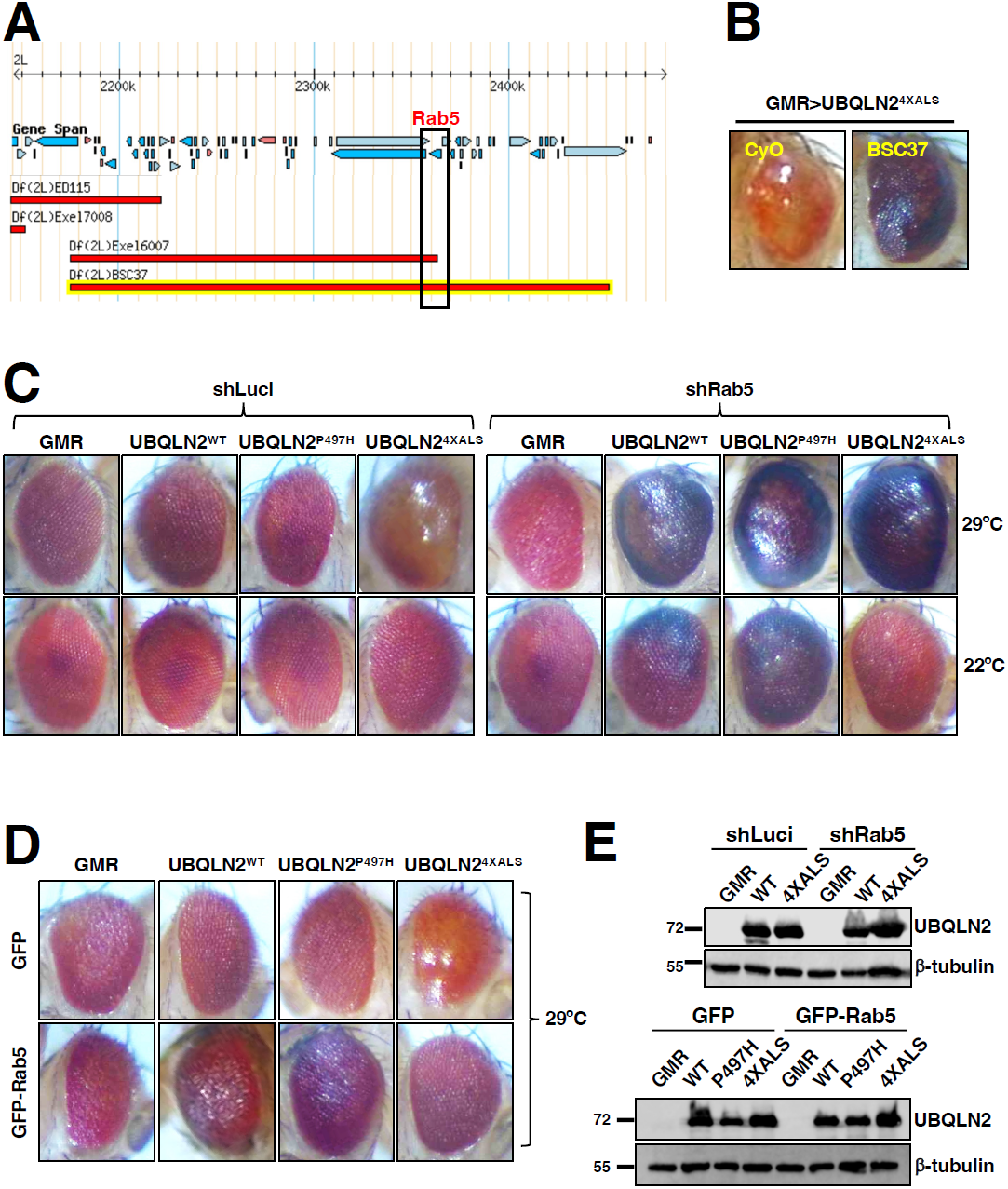
*Rab5* is a UBQLN2^4XALS^ modifier gene. **(A)** Genomic map of BSC37. The region spanning the *Rab5* gene is boxed. **(B)** Representative eye images from GMR>UBQLN2^4XALS^/BSC37 and GMR>UBQLN2^4XALS^/CyO flies. **(C)** Differential effects of Rab5 knockdown on GMR>UBQLN2 eye phenotypes at 22°C and 29°C. Recombinant GMR>UBQLN2 flies of indicated genotypes were crossed to flies expressing shRNAs targeting Rab5 (shRab5) or luciferase (shLuci). F1 progeny were processed for eye imaging 2-3 days after eclosion. **(D)** GFP-Rab5 overexpression rescued the UBQLN2^4XALS^ RE phenotype at 29°C. GMR>UBQLN2 flies of indicated genotypes were crossed to flies harboring UAS-GFP or UAS-GFP-Rab5 transgenes. **(E)** UBQLN2 expression levels in whole heads of GMR-Gal4, GMR>UBQLN2^WT^, GMR>UBQLN2^P497H^, or GMR>UBQLN2^4XALS^ flies on the indicated genetic backgrounds. Neither Rab5 knockdown, nor Rab5 overexpression appreciably impacted UBQLN2 expression.

A similar approach was used to map the causal UBQLN2^4XALS^ suppressor gene in BSC180. Two different overlapping Dfs (C144 and ED4651) rescued UBQLN2^4XALS^ eye toxicity to a similar extent as BSC180 (Figure supplement 3A, B). Within this overlap we identified lilliputian (*lilli*) as a gene of interest. *lilli* encodes a transcriptional elongation factor that is orthologous to mammalian AF4/FMR2 (fragile X mental retardation 2)^34^. Interestingly, *lilli* mutations were recently shown to suppress toxicity of TDP-43 and *C9ORF72*-derived DPRs in *Drosophila*^35,36^. In support of *lilli* as a UBQLN2^4XALS^ suppressor, a *lilli*^*A1*7-2^ LOF allele diminished UBQLN2^4XALS^-mediated eye toxicity to a similar extent as *lilli*-spanning Dfs (Figure supplement 3B). *lilli* may be of interest as a general suppressor of toxicity arising from misexpression of ALS-associated genes in Drosophila, but was not further investigated in this study.

### Unc-5 mutations suppress UBQLN2 toxicity

Next, we mapped a UBQLN2^4XALS^ suppressor in the genetic interval spanned by ED2426 and BSC346, which suppressed GMR>UBQLN2^4XALS^ eye phenotypes to a similar extent (Figure 4A, B). The region of overlap between ED2426 and BSC346 contains *Unc-5*, which encodes a transmembrane dependence receptor that mediates axonal repulsion and apoptosis suppression in response to secreted netrin ligands^37-42^.

**Figure 4.**
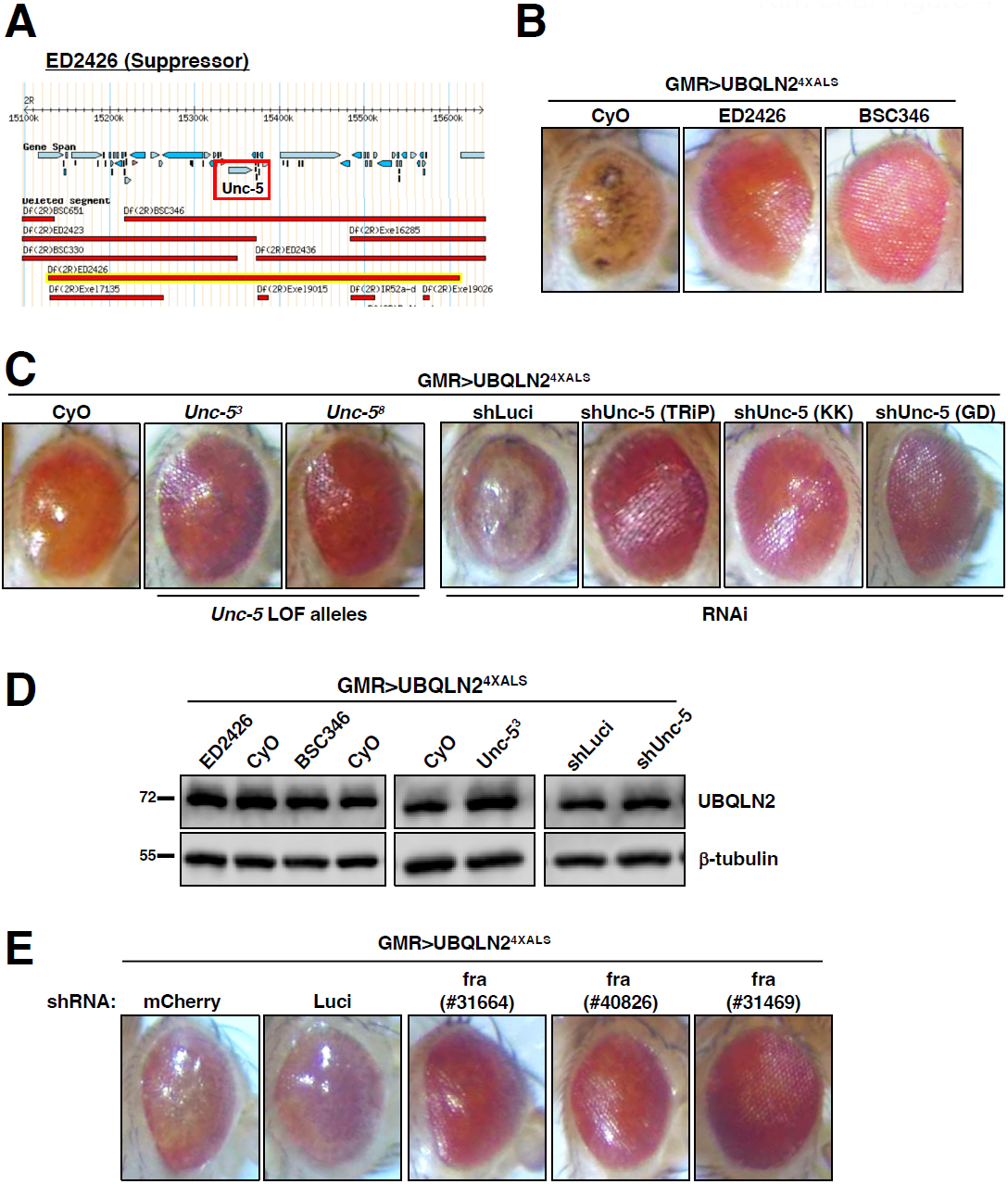
Disruption of *Unc-5* suppresses UBQLN2-associated eye degeneration. **(A)** Schematic depiction of ED2426 and overlapping Dfs in relation to the *Unc-5* gene locus. **(B)** Representative eye images of GMR>UBQLN2^4XALS^ flies crossed to Df lines ED2426 and BSC346. **(C)** Single copy expression of *Unc-5* LOF alleles (left panels) or three independent Unc-5 RNAi alleles diminished the RE phenotype of GMR>UBQLN2^4XALS^ flies at 29°C. **(D)** UBQLN2 expression levels in whole heads of GMR>UBQLN2^4XALS^ flies on the indicated genetic backgrounds. **(E)** Fra silencing reduced the RE phenotype of GMR>UBQLN2^4XALS^ flies. Shown are representative ye phenotypes of F1 progeny from GMR>UBQLN2^4XALS^ flies crossed to control (shLuci, shmCherry), or Fra RNAi lines at 29°C.

In support of a genetic interaction between *Unc-5* and UBQLN2^4XALS^, two *Unc-5* LOF alleles^38^, and three different RNAi lines diminished the RE phenotype of GMR>UBQLN2^4XALS^ flies without affecting UBQLN2 expression (Figure 4C, D). By contrast, Unc-5 knockdown had no effect on the RE phenotype caused by mutant FUS expression (Figure supplement 4). The repulsive activity of Unc-5 on axon guidance is antagonized by *frazzled (fra)*, which encodes the fly ortholog of mammalian deleted in colon carcinoma (DCC)^43-45^. *fra* silencing by two different RNAi lines rescued the UBQLN2^4XALS^ eye phenotype to a similar extent as *Unc-5* knockdown (Figure 4E). The combined findings suggest signaling through Unc-5/Frazzled potentiates UBQLN2^4XALS^ toxicity in the Drosophila compound eye.

To evaluate the impact of Unc-5 silencing on motor function, we measured climbing behavior of recombinant flies expressing UBQLN2^4XALS^ under control of a D42 motor neuron driver in the presence of shUnc-5 or control shRNAs at 29°C. The moderate climbing defect of D42>UBQLN2^4XALS^ flies relative to D42-Gal4 flies was partially reversed by two different shUnc-5 alleles in comparison to shLuci and shmCherry controls (Figure 5A). Consistent with climbing defects, UBQLN2^4XALS^ caused neuromuscular junction (NMJ) abnormalities, including increased numbers of satellite boutons and reduced numbers of mature boutons. Both phenotypes were corrected by Unc-5 silencing (Figure 5B, C). Finally, we also measured the effect of Unc-5 silencing on lifespans of flies expressing UBQLN2^4XALS^ under control of a pan-neuronal Elav driver. Elav>UBQLN2^4XALS^/shUnc-5 flies showed a significant increase in lifespan relative to Elav>UBQLN2^4XALS^ flies crossed to shLuci, with the effect being most pronounced in female flies (Figure supplement 5A). Unc-5 knockdown also modestly increased the lifespan of male Elav-Gal4 flies while having no impact on lifespan of Elav-Gal4 female flies (Figure supplement 5B). Altogether, these experiments suggest that aberrant Unc-5 signaling contributes to neuronal phenotypes in UBQLN2^4XALS^ flies.

**Figure 5.**
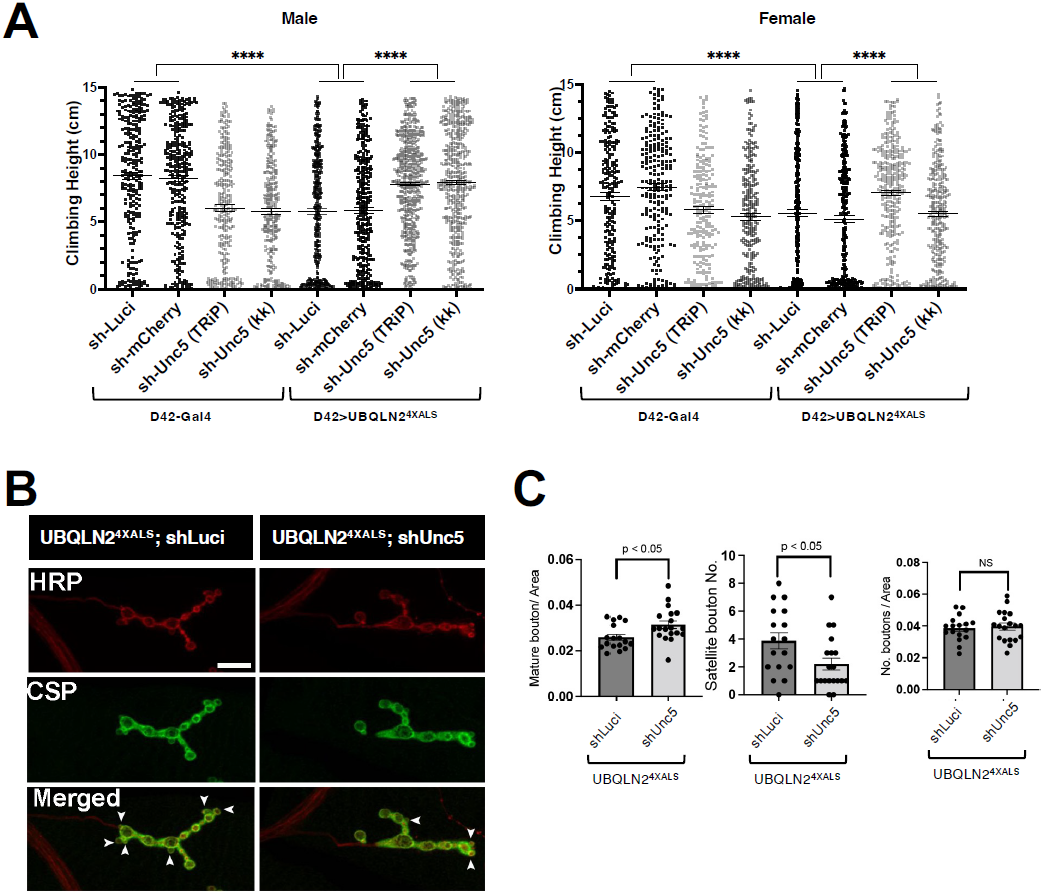
*Unc-5* silencing in iMNs suppressed UBQLN2^4XALS^-associated climbing defects. **(A)** Recombinant D42>UBQLN2^4XALS^ flies expressing UBQLN2 under control of the motor neuron-specific D42 driver were crossed to the indicated RNAi lines. Climbing potential of male (left) and female (right) progeny reared at 29°C was measured 7-days after eclosion as described in Methods. Data analysis was performed using ordinary one-way ANOVA. Data are shown as mean +/- SEM. n>100 flies, \*\*\*\**p* ≤ 0.0001. **(B)** NMJ morphology analysis of D42 > UBQLN2^4XALS^ larvae expressing the indicated shRNAs. NMJs dissected from 3^rd^ instar larvae were stained with α-HRP and α-CSP antibodies and imaged by confocal microscopy. **(C)** Number of NMJs harboring indicated phenotypes were tabulated from greater than 50 NMJs per genotype. Unpaired *t*-test was used for statistical analysis. Data are shown as mean +/- SEM.

### The motor neuron guidance factor *beat-1b* suppresses UBQLN2^4XALS^ eye toxicity

The identification of Unc-5 as a UBQLN2^4XALS^ suppressor raised the possibility that axonal guidance defects are particularly relevant to the UBQLN2 toxicity mechanism. Interestingly, the overlapping genetic interval spanned by the UBQLN2^4XALS^ suppressors Exel6038 and r10 contains *beat-1b* and *beat-1c*, two members of the *beaten path* (*beat*) family of axon guidance genes (Figure supplement 6A-C)^46^. Neuronally-expressed Beat proteins regulate motor axon guidance and defasciculation in response to Sidestep (Side) ligands expressed on target substrates^47-50^. UBQLN2^4XALS^ flies crossed to an shRNA line targeting *beat-1b* or a line harboring a P-element insertion in the *beat-1b* coding sequence exhibited less severe eye degeneration than UBQLN2^4XALS^ flies crossed to control RNAi lines (Figure supplement 6D, E). By contrast, a *beat-1c* RNAi allele did not rescue the GMR>UBQLN2^4XALS^ RE phenotype (Figure supplement 6D). These findings support the idea that Beat signaling contributes to UBQLN2 toxicity and further implicate axonal guidance defects as a contributing pathomechanism to UBQLN2-associated neurodegeneration.

### iPSC models for UBQLN2-associated ALS

We employed CRISPR/CAS9 to introduce ALS-associated mutations into the X-linked *UBQLN2* gene in neonatal male iPSCs (WC031i-5907-6, see Methods and Ref.^51^). During the course of this work, we serendipitously derived an *UBQLN2* allele harboring clinical P497H and P525S mutations (termed UBQLN2^2XALS^) as well as a UBQLN2^I498X^ allele that truncated the UBQLN2 ORF at codon 498 within the PRR. The expression and RIPA solubility of UBQLN2^P497H^, UBQLN2^2XALS^, and UBQLN2^4XALS^ were comparable to UBQLN2^WT^ in undifferentiated iPSCs, while UBQLN2^I498X^ could not be detected by Western blotting suggesting it is a null allele (Figure 6A). We also failed to detect cytologic UBQLN2^P497H^, UBQLN2^2XALS^, or UBQLN2^4XALS^ aggregates in immunostaining experiments, indicating that endogenous ALS mutations are insufficient to promote UBQLN2 aggregation in undifferentiated iPSCs (Figure 6B). As expected, UBQLN2^I498X^ iPSCs exhibited very weak immunoreactivity with UBQLN2 antibodies.

**Figure 6.**
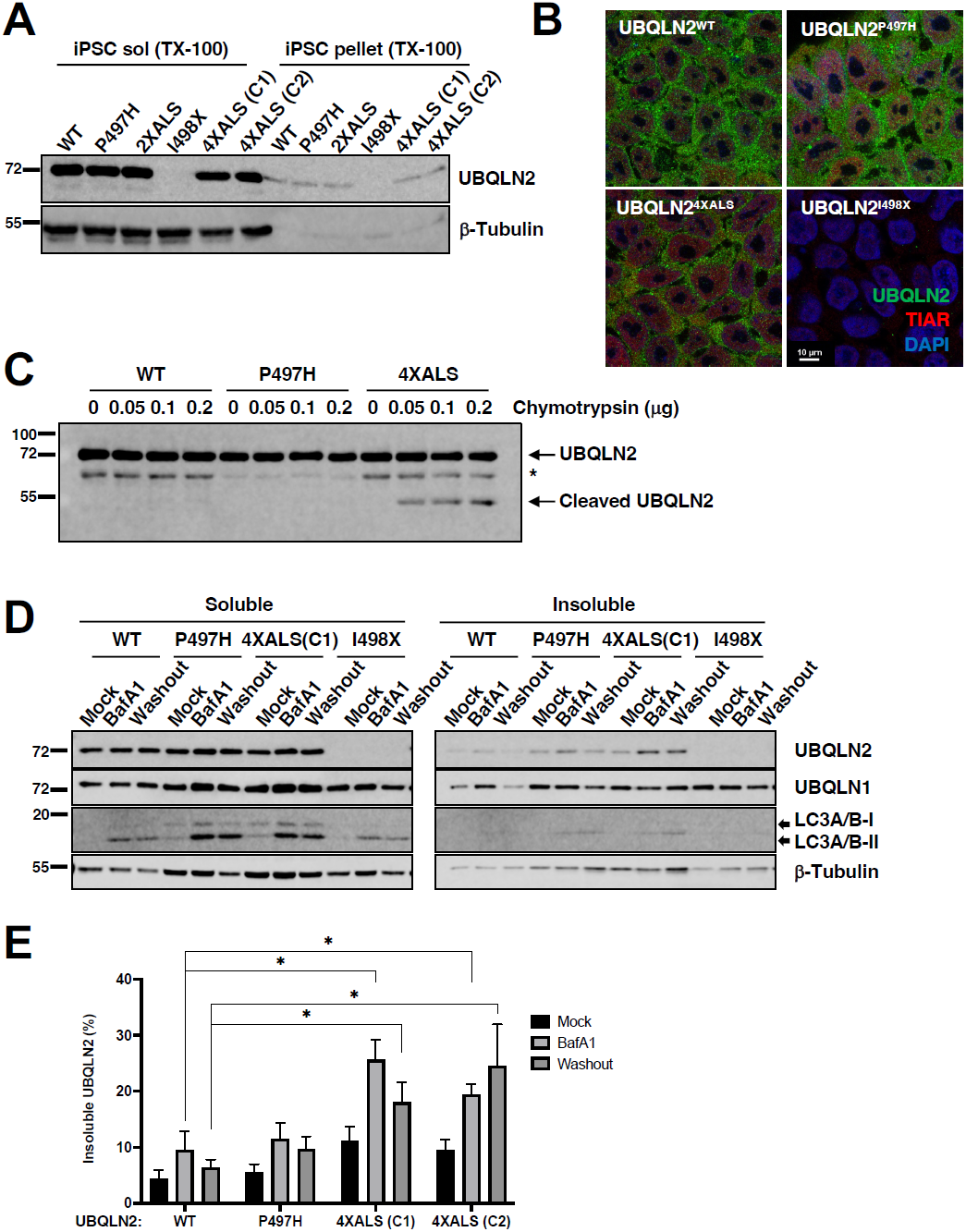
Localization and solubility of UBQLN2^ALS^ mutants in iPSCs. **(A)** Extracts from UBQLN2^WT^, UBQLN2^P497H^, UBQLN2^2XALS^, UBQLN2^I498X^, or UBQLN2^4XALS^ iPSCs (clone 1(C1) and clone 2 (C2)) were separated into soluble and insoluble fractions in 1% Triton X-100 (TX-100) buffer and immunoblotted with α-UBQLN2, and α-β-tubulin antibodies. **(B)** Localization patterns of wild-type and UBQLN2^ALS^ proteins in iPSCs. UBQLN2^WT^, UBQLN2^P497H^, UBQLN2^4XALS^, and UBQLN2^I498X^ iPSCs were stained with α-UBQLN2 and α-TIAR antibodies and imaged by confocal microscopy. Note the lack of cytosolic aggregates. **(C)** Cell extracts from iPSCs of the indicated genotypes were incubated at room temperature with increasing amounts of chymotrypsin for 5 min. After separation by SDS-PAGE, the proteins were immunoblotted with α-UBQLN2 antibodies. Positions of full-length and cleaved UBQLN2 are denoted by arrows. *: non-specific band. **(D)** Autophagy inhibition with BafA1 reduced solubility of endogenous UBQLN2^ALS^ proteins. UBQLN2^WT^, UBQLN2^P497H^, UBQLN2^4XALS^, and UBQLN2^I498X^ iPSCs were treated with 100 nM of BafA1 for 16 h followed by BafA1 washout and incubation in BafA1-free growth media for 8 h. Detergent extracts were separated into soluble and insoluble fractions and analyzed by SDS-PAGE and immunoblotting using UBQLN2, UBQLN1, LC3A/B, and b-tubulin antibodies. **(E)** Quantification of detergent soluble and insoluble UBQLN2 following BafA1 treatment. Unpaired *t*-test was used for statistical analysis. Data are shown as mean +/- SEM. n=3, \**p* ≤ 0.05.

Previous work demonstrated that the 4XALS mutation increased the chymotrypsin sensitivity of purified UBQLN2, likely due to defective folding of the PRR^31^. To determine whether the 4XALS mutation altered the folding of endogenous UBQLN2, we incubated detergent extracts from UBQLN2^WT^, UBQLN2^P497H^, UBQLN2^4XALS^ iPSCs with increasing concentrations of chymotrypsin. As shown in Figure 6C, UBQLN2^4XALS^ exhibited a unique chymotryptic fragmentation pattern relative to UBQLN2^WT^ and UBQLN2^P497H^, suggesting that endogenous UBQLN2^4XALS^ is misfolded.

Reasoning that transient inhibition of protein degradation may potentiate UBQLN2 aggregation, we prepared soluble and insoluble fractions of UBQLN2^WT^, UBQLN2^P497H^, and UBQLN2^4XALS^ iPSCs exposed to the proteasome inhibitor MG132 or the autophagy inhibitor bafilomycin A1 (BafA1). While MG132 had no effect (not shown), BafA1 decreased the solubility of both wild-type and mutant UBQLN2 proteins, with the effect being most pronounced for UBQLN2^4XALS^ (Figure 6D).

In addition, while the solubility of UBQLN2^WT^ and UBQLN2^P497H^ recovered following BafA1 washout, the fraction of insoluble UBQLN2^4XALS^ remained elevated (Figure 6E). Levels of LC3 lipidation did not appreciably differ between the four genotypes in the absence or presence of BafA1 (Figure 6D). Upon BafA1 treatment, iPSCs of all three UBQLN2 genotypes exhibited cytosolic punctae that were partially localized with the lysosomal marker LAMP1 (Figure 7A). By contrast, UBQLN2 punctae did not significantly overlap with Rab5-positive endosomes (Figure supplement 7). These findings suggest that UBQLN2^ALS^ mutants exhibit prolonged insolubility upon inhibition of autophagy.

**Figure 7.**
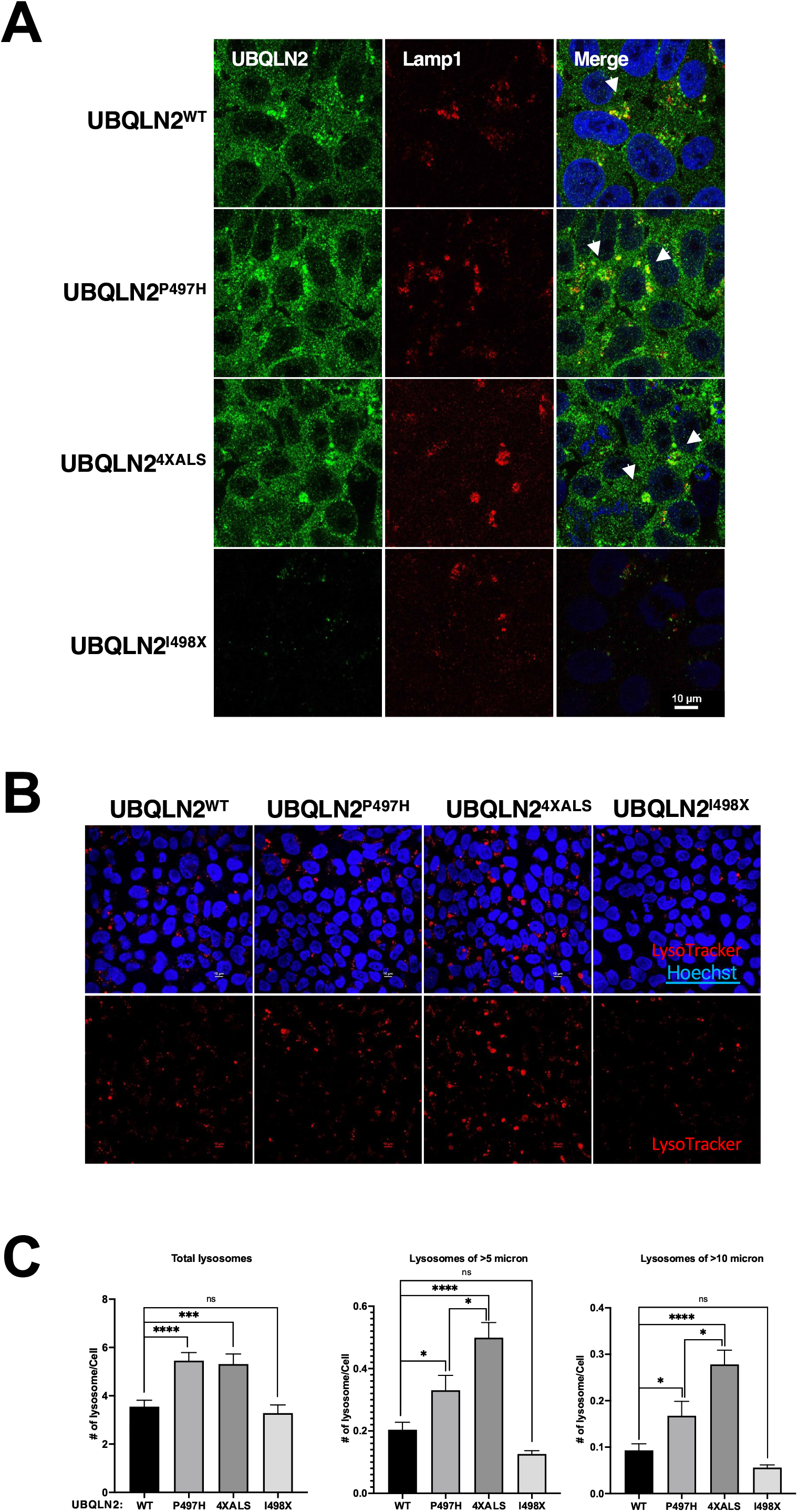
Endogenous UBQLN2^ALS^ mutants perturb lysosomes. **(A)** UBQLN2^WT^, UBQLN2^P497H^, UBQLN2^4XALS^, and UBQLN2^I498X^ iPSCs were treated with 100 nM of BafA1 for 16 h and processed for immunostaining with UBQLN2 and Lamp1 antibodies. Arrowheads indicate colocalization of UBQLN2 and Lamp1. **(B, C)**Increased lysosomal size and number in UBQLN2^ALS^ mutant iPSCs. **(B)** Representative images from UBQLN2^WT^, UBQLN2^P497H^, UBQLN2^4XALS^, and UBQLN2 ^I498X^ iPSCs following labeling with LysoTracker™ Red DND-99 for 1 h and costaining with Hoechst 33342. Scale bar: 10 μm. **(C)** Size and numbers of LysoTracker-positive compartments were analyzed on a per cell basis using Fiji. Unpaired *t*-test was used for statistical analysis. Data are shown as mean +/- SEM. \**p* ≤ 0.05, \*\*\**p* ≤ 0.001, \*\*\*\**p* ≤ 0.0001.

Given colocalization of UBQLN2 with LAMP1, we evaluated impacts of P497H and 4XALS mutations on lysosomal number and size in live cells using the lysosome-tropic fluorescent probe, LysoTracker. Both the abundance and average size of lysosomes were significantly elevated in UBQLN2^P497H^ and UBQLN2^4XALS^ iPSCs, relative to UBQLN2^WT^ or UBQLN2^I498X^ iPSCs, which exhibited qualitatively similar LysoTracker staining patterns (Figure 7B). In particular, the frequency of lysosomes greater than 5 μm and 10 μm was significantly elevated in UBQLN2^4XALS^ iPSCs versus UBQLN2^P497H^ iPSCs (Figure 7C).

### UBQLN2^ALS^ iMNs exhibit axonal inclusions and neurite defects

A failure to detect UBQLN2^ALS^ aggregates in untreated iPSCs could be due to continuous cytosolic dilution of UBQLN2 during mitotic cell division or may reflect the absence of neuron-specific stimuli that promote UBQLN2 aggregation. To explore these ideas, we carried out immunostaining using iMNs expressing UBQLN2^WT^, UBQLN2^P497H^, UBQLN2^2XALS^, or UBQLN2^4XALS^ (Figure 8A and data not shown). UBQLN2^WT^, UBQLN2^P497H^, and UBQLN2^2XALS^ iMNs showed largely diffuse localization patterns, whereas UBQLN2^4XALS^ formed discrete aggregates distributed throughout iMN cell bodies and axons (Figure 8A). F-actin labeling with phalloidin revealed that UBQLN2^WT^ and UBQLN2^P497H^ were evenly distributed throughout growth cone lamellipodia and filopodial spikes. By contrast, lamellipodial UBQLN2^4XALS^ staining was generally weaker, with the exception of occasional brightly-staining aggregates that were observed in ∼50% of growth cones examined (Figure 8B).

**Figure 8.**
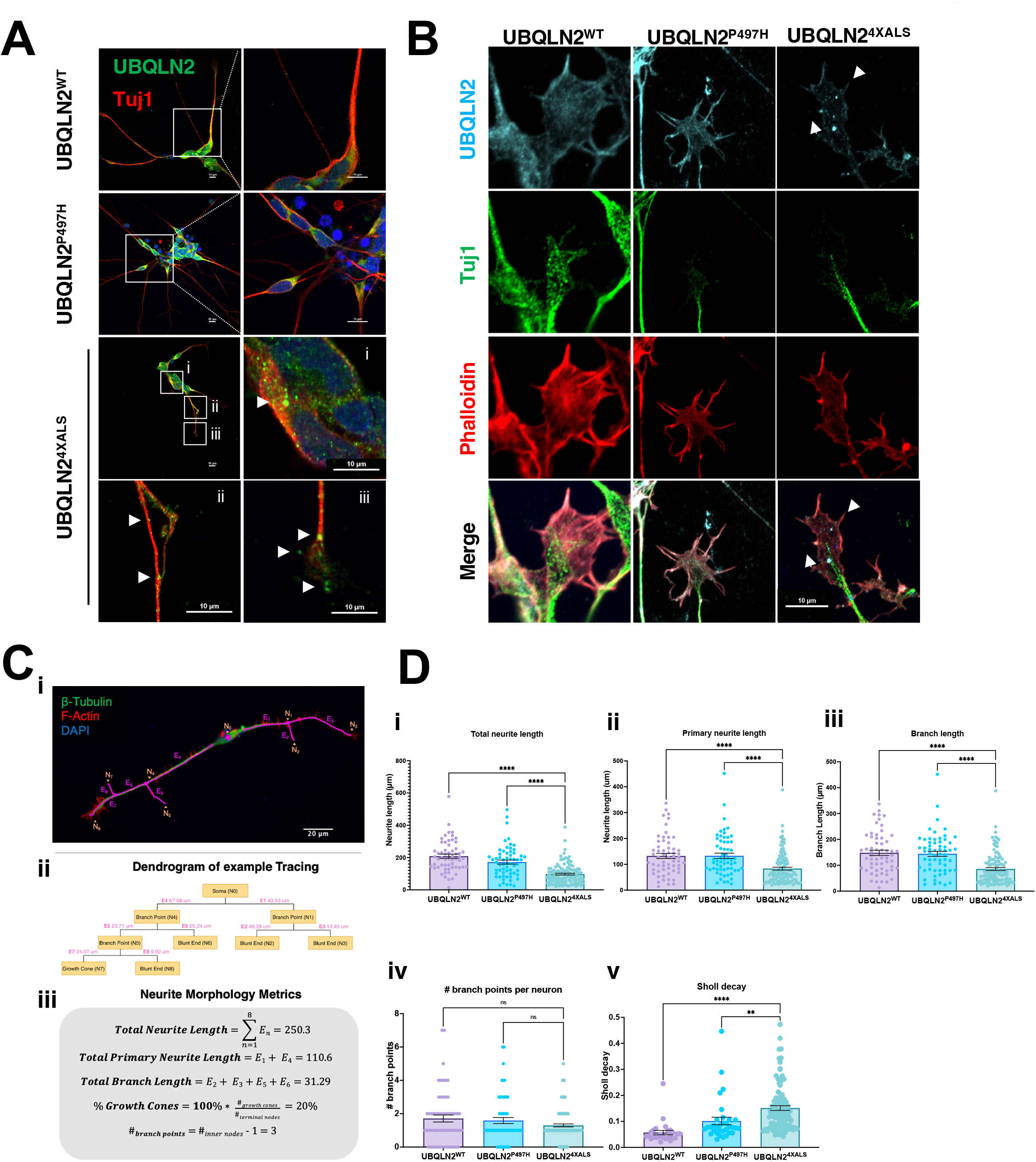
Protein aggregation and neurite defects in UBQLN2^ALS^ iMNs. **(A)** UBQLN2^WT^, UBQLN2^P497H^, and UBQLN2^4XALS^ iMNs were immunostained for UBQLN2 and Tuj1. Magnified images of UBQLN2^4XALS^ iMN soma (i), axons (ii), and neurite terminals (iii) are shown; aggregates are marked with arrows. **(B)** UBQLN2 localizes to growth cone lamellipodia and filopodia. Differentiated iMNs of the indicated genotypes were costained for UBQLN2, Tuj1, and filamentous actin (phalloidin). Note reduced complexity of the UBQLN2^4XALS^ growth cone. Arrowheads indicate UBQLN2^4XALS^ aggregates. Scale bars = 10 μm. **(C)** Schematic of Sholl analysis. Tracing example of an iMN (i) displays the paths representing individual neuron structure, with nodes (N_i_) and edges (E_j_) corresponding to the dendrogram on (ii) and (iii). **(D)** UBQLN2^4XALS^ iMNs exhibit reduced complexity. UBQLN2^WT^, UBQLN2^P497H^, and UBQLN2^4XALS^ iMNs were stained with a-Tuj1 and imaged by confocal microscopy. One hundred neurons of the indicated genotypes were traced using Simple Neurite Tracer (SNT) and subjected to Sholl image analysis to quantify total neurite projection path length (i), primary neurite length (ii), terminal neurite length (iii), neurite branch points (iv), and Sholl decay (v). Data analysis was performed using ordinary one-way ANOVA. Data are shown as mean +/- SEM. n>100 iMNs, \*\**p* ≤ 0.01, \*\*\*\**p* ≤ 0.0001.

Neurite defects in UBQLN2^4XALS^ iMNs were quantified using Sholl analyses according to the diagram shown in Equation 1 and Figure 8C, respectively. UBQLN2^4XALS^ iMNs exhibited a significant reduction in average total neurite length, primary neurite length, and neurite branch length relative to UBQLN2^WT^ iMNs, which manifested as an increased rate of Sholl decay (Figure 8D). By contrast, neurite length and complexity were comparable between UBQLN2^P497H^ and UBQLN2^WT^ iMNs, indicating the clinical P497H mutation is insufficient to disrupt neurite growth dynamics (Figure 8D). Aggregates and neurite defects were also observed using UBQLN2^4XALS^ iMNs differentiated from an independently derived UBQLN2^4XALS^ iPSC line, suggesting they are a specific consequence of the 4XALS mutation (Figure supplement 8A, B).

### UNC5B and DCC silencing reduce neurite and growth cone defects in UBQLN2^4XALS^ iMNs

Mammals harbor a single *fra* ortholog, *DCC*, and four closely related paralogs with homology to *Unc-5*: *UNC5A, UNC5B, UNC5C, and UNC5D*. Among these, we focused on *UNC5B*, which is widely expressed in nervous tissue and has well-described roles in axon guidance and apoptosis regulation^41,42,52,53^. To assess contributions of DCC and UNC5B signaling to UBQLN2-associated toxicity, we transduced UBQLN2^4XALS^ iPSCs with lentiviral shRNA vectors targeting DCC or UNC5B and differentiated the cells into iMNs for neurite analysis. qPCR confirmed that expression of UNC5B and DCC was reduced ∼40-60% in their respective shRNA-transduced iPSCs relative to iPSCs transduced with a non-targeting shRNA vector (Figure supplement 9A, B). UBQLN2^4XALS^ iMNs expressing DCC shRNA also showed reduced DCC immunoreactivity at filopodial spikes (Figure supplement 9C). Both UNC5B and DCC knockdown significantly increased average total neurite length, primary neurite length, branch length, and branch point abundance in UBQLN2^4XALS^ iMNs relative to UBQLN2^4XALS^ iMNs expressing a non-targeting (NT) control shRNA (Figure 9A).

**Figure 9.**
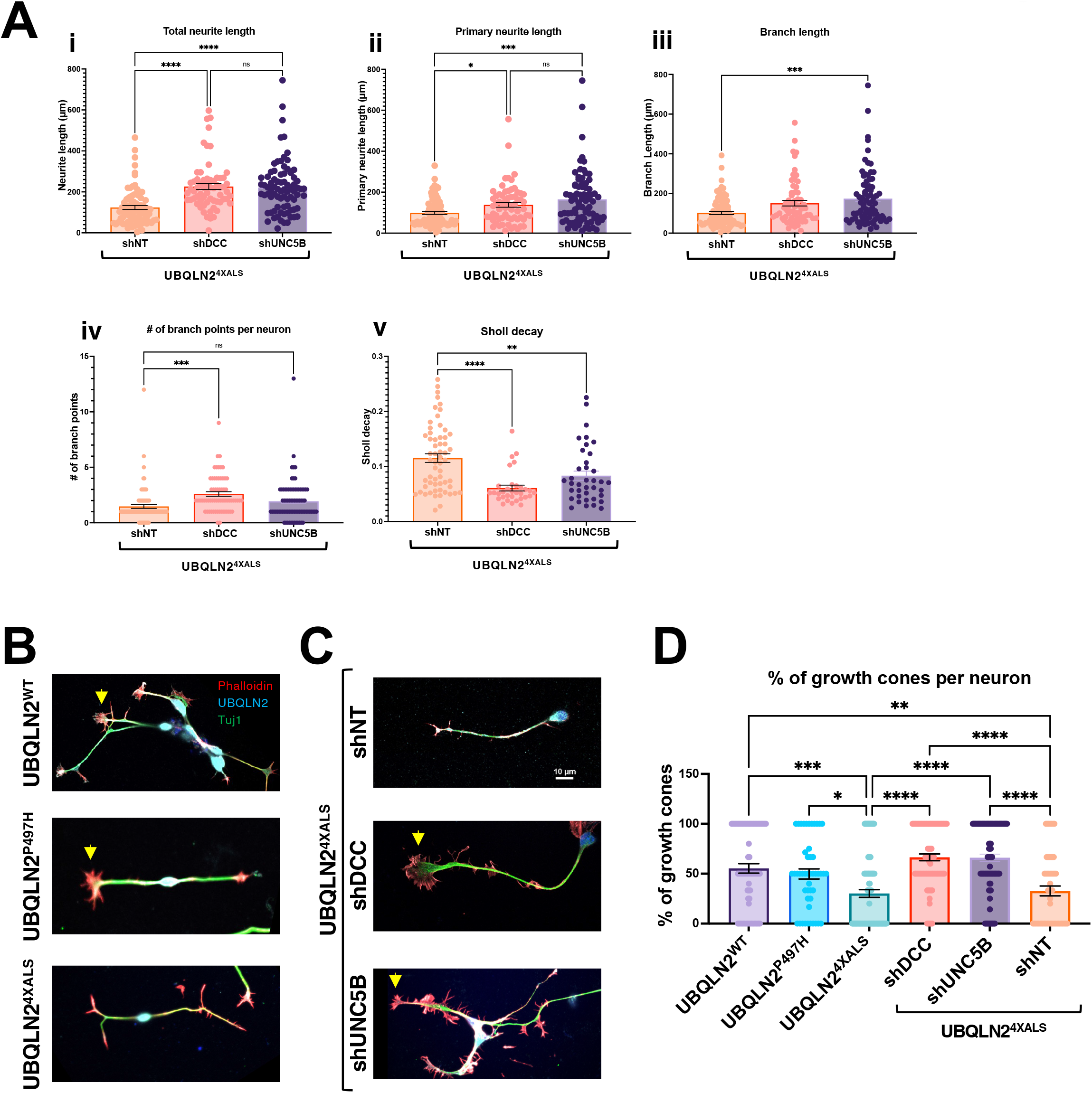
UNC5B and DCC silencing partially reverse neurite and growth cone defects in UBQLN2^4XALS^ iMNs. **(A)** UBQLN2^4XALS^ iMNs expressing shNT, shUNC5B, or shDCC were subjected to Sholl analysis of neurite length and complexity as described in Figure 8. Data analysis was performed using ordinary one-way ANOVA. Data are shown as mean +/- SEM. n>100 iMNs, \**p* ≤ 0.05, \*\**p* ≤ 0.01, \*\*\**p* ≤ 0.001, \*\*\*\**p* ≤ 0.0001. **(B)** Growth cone morphologies in UBQLN2^4XALS^ iMNs. UBQLN2^WT^, UBQLN2^P497H^, and UBQLN2^4XALS^ iMNs. iMNs of the indicated genotypes were stained with a-UBQLN2, a-Tuj1, and phalloidin. Note reduced growth cone elaboration in UBQLN2^4XALS^ iMNs. **(C)** Enhanced growth cone elaboration in UBQLN2^4XALS^ iMNs expressing DCC or UNC5B shRNAs. iMNs of the indicated genotype were stained with a-UBQLN2, a-Tuj1, and phalloidin. Arrowheads denote growth cones. **(D)** Quantification of growth cones in UBQLN2^WT^, UBQLN2^P497H^, and UBQLN2^4XALS^, and UBQLN2^4XALS^ iMNs expressing the indicated shRNAs. Data analysis was performed using ordinary one-way ANOVA. Data are shown as mean +/- SEM. n>100 iMNs, \**p* ≤ 0.05, \*\**p* ≤ 0.01, \*\*\**p* ≤ 0.001, \*\*\*\**p* ≤ 0.0001.

We next evaluated growth cone morphology in wild-type and UBQLN2^ALS^ iMNs stained with phalloidin. Compared to UBQLN2^WT^ or UBQLN2^P497H^ iMNs, UBQLN2^4XALS^ iMNs showed a high proportion of blunt-end termini versus growth cone termini, suggesting a defect in growth cone elaboration (Figure 9B, D). By contrast, UBQLN2^4XALS^ iMNs expressing UNC5B or DCC shRNAs exhibited supernumerary growth cones along primary and secondary neurites and an increase in growth cone size and relative abundance relative to blunt-end termini (Figure 9C, D). These findings suggest that aberrant DCC-UNC5 signaling suppresses growth cone elaboration in UBQLN2^4XALS^ iMNs and that axon guidance defects contribute to toxicity phenotypes in fly and iMNs models for UBQLN2-associated ALS.

## Discussion

In this study we investigated how ALS-associated mutations in the Ub chaperone UBQLN2 cause cellular toxicity in Drosophila and gene-edited iMNs. Our findings support roles for endolysosomal dysfunction and axonal guidance defects as disease drivers in UBQLN2-associated ALS.

A screen of 194 Dfs against UBQLN2^P497H^ and UBQLN2^4XALS^ alleles with differing aggregation potential and toxicity spectra resulted in the identification of 35 Dfs (7 suppressors and 28 enhancers) that influenced eye toxicity. We successfully mapped three of the four non-overlapping UBQLN2^4XALS^ suppressor loci to implicate *Unc-5, lilli*, and *beat-1b* as UBQLN2^4XALS^ modifier genes. Overlapping Dfs failed to phenocopy the strong suppressor phenotype of BSC19 and a candidate approach failed to identify a causal gene in the BSC19 interval.

Among the 28 enhancer loci, we successfully mapped *Rab5* as the causal gene in BSC37. Rab5 silencing caused a hyperpigmented eye phenotype in flies expressing either wild-type or ALS-mutant UBQLN2 alleles, suggesting that overexpressed UBQLN2 proteins interfere with early endosomal function. This finding is congruent with recent work by Senturk et al. describing a role for endogenous *dUbqln* in endolysosomal acidification^54^. While we attempted to map causal genes in several other UBQLN2 enhancer loci, we were unable to identify candidate genes whose silencing fully replicated the degenerative eye phenotypes seen with the Df crosses. A plausible explanation for this is that the disruption of multiple genes is responsible for the enhancer effects of some Df lines. Nevertheless, other UBQLN2 modifier genes await discovery.

The UBQLN2^4XALS^ suppressor *lilli* (Figure supplement 3) was also identified as a phenotypic suppressor in Drosophila overexpression models for TDP-43 and poly(GR) dipeptide repeats (DPRs) linked to C9ORF72-associcated ALS^36,55^. *lilli* is an orthologue of mammalian FMR2 (fragile X mental retardation 2), which plays a role in transcriptional elongation. lilli/FMR2 silencing was shown to selectively reduce transcription of G4C2 HREs in Drosophila and iPSC-derived neurons^36,55^. It is conceivable that Lilli is similarly required for read-through of the GC-rich UBQLN2 ORF. Alternatively, because *lilli* was also identified as a suppressor of Hairless overexpression toxicity in the fly eye^56^, *lilli* LOF alleles may suppress overlapping death pathways engaged by neurodegeneration-associated proteins.

Our findings suggest that axon guidance pathways play an important role in UBQLN2-mediated toxicity. Unc-5 fulfills dual functions as a repulsive axon guidance factor and neuronal dependence receptor, and either or both functions could be relevant to the phenotypic rescue conferred by Unc-5 silencing in flies expressing the UBQLN2^4XALS^ transgene^37-39^. Knockdown of Fra, which mediates chemoattractive responses to netrin, also partially rescued UBQLN2^4XALS^-associated eye and motor neuron phenotypes (Figure 4E, Figure 9A, C, D). One interpretation of this data is that heterodimeric Unc-5-Fra complexes mediate toxicity initiated by UBQLN2^ALS^ mutants. Alternatively, Unc-5 and Fra/DCC may function in partially redundant fashion to instigate toxicity in UBQLN2^ALS^ flies.

The identification of *beat-1b* as a UBQLN2 modifier further supports axon pathfinding defects as a disease driver in the UBQLN2^4XALS^ flies. Beat proteins mediate guidance of motor neuron axons through transient interactions with transmembrane Sidestep receptors whose expression pattern on muscle, muscle stem cells, and neurons constitutes a stereotypic guidance path^47,49,57,58,59^. Although no clear human ortholog exists, Beats exhibit weak homology to mammalian NekL3/SynCAM2/CADM2, a nectin-like molecule that mediates adhesion of myelinated axons and oligodendrocytes^60-62^.

We developed UBQLN2 gene-edited iPSCs and iMNs to investigate UBQLN2 pathomechanisms at the cellular level. Endogenous UBQLN2^P497H^ and UBQLN2^4XALS^ exhibited wild-type localization and solubility in iPSCs in the absence of stress; however, the relative insolubility of UBQLN2^4XALS^ seen in overexpression studies was unmasked when iPSCs were treated with the lysosomal acidification inhibitor BafA1 (Figure 6D) or following neuronal differentiation (Figure 8A, B). UBQLN2^4XALS^ aggregates were distributed throughout the soma, axon, and growth cone lamellipodia extending into filopodial tips, raising the possibility that such aggregates interfere with growth cone dynamics. Indeed, UBQLN2^4XALS^ iMNs exhibited reduced neurite length, diminished neurite complexity, and reduced growth cone numbers relative to wild-type and UBQLN2^P497H^ iMNs (Figure 8D, 9B, 9D). While these findings imply that UBQLN2^4XALS^ toxicity is tightly linked to its aggregation potential, it remains possible that soluble forms of UBQLN2^4XALS^ and clinical UBQLN2^ALS^ proteins also disrupt key cellular processes. Given genetic interactions between UBQLN2 and Rab5 in flies (Figure 3) and the lysosomal defects seen in UBQLN2^P497H^ and UBQLN2 iPSCs (Figure 6), the endolysosomal pathway may be a particularly relevant site of UBQLN2 toxicity.

Knockdown of UNC5B and DCC significantly increased neurite length and complexity and partially corrected growth cone morphology defects of UBQLN2^4XALS^ iMNs, indicating that the UBQLN2^4XALS^ toxicity mechanism is at least partially conserved between flies and humans (Figure 9). As in Drosophila, mammalian DCC/Fra and UNC5 family receptors have been implicated in both axon guidance and apoptosis, and either or both of these biological activities may be relevant to their genetic interactions with UBQLN2^37,40-42,53,63,64^. UNC5 and DCC/Fra harbor extended cytoplasmic domains that regulate caspase activation and are themselves targets for caspase mediated cleavage^41,65^. Engagement by netrin ligand is thought to suppress the intrinsic apoptotic potential of the UNC5 death domain while γ-secretase mediated cleavage of UNC5C has been linked to neuronal apoptosis in AD^41,52,62^. The contributions of UNC5B death signaling to toxicity phenotypes in UBQLN2^ALS^ iMNs and flies may be informed by separation of function mutants with reduced caspase recruitment potential. In addition, the contributions of other UNC5 paralogs to UBQLN2 toxicity remains to be determined.

Distal axon and synaptic defects are implicated in ALS pathogenesis and have been observed in diverse ALS disease models. For instance, reduced expression of the microtubule binding protein Stathmin 2 due to misregulation of its RNA splicing and/or polyadenylation is strongly linked to axonal growth and regeneration defects in TDP-43 associated ALS^66,67^. Single nucleotide polymorphisms that promote inclusion of a cryptic cassette exon in the presynaptic regulator *UNC13A* are a risk factor for fALS^68^, whereas loss of nuclear TDP-43 has been linked to UNC13A missplicing and synaptic defects in sALS^69,70^. While *UNC5* has not been implicated as an ALS gene, signaling downstream of UNC5 may contribute to axonal retraction and/or synaptic phenotypes initiated by UBQLN2 mutation, TDP-43 aggregation, or mutations in ALS-associated genes.

Finally, while findings in flies and iMNs support a broad role for axon guidance defects in the UBQLN2 toxicity mechanism, there are several limitations to our study. First, despite attempts to focus on mutation-dependent suppressors, overexpression of wild-type and ALS-mutant UBQLN proteins can cause disease non-specific toxicities that may confound phenotypic screens in flies. Second, the UBQLN2^4XALS^ mutant, while enabling phenotypic screens, is not a bona fide disease allele and may elicit toxicities unrelated to those caused by clinical ALS mutations. Finally, neonatal iPSCs and their derivative iMNs, while possessing numerous strengths, are unlikely to capture age-dependent abnormalities that contribute to neurodegeneration in an intact human nervous system.

## Materials and methods

### Drosophila methods

Flies were maintained with the standard cornmeal-yeast medium (Nutri-Fly BF #66-112, Genesee scientific) supplemented with propionic acid and all crosses were performed at 22°C. For heat stress experiments, all crosses were performed at indicated temperatures (27°C or 29°C). Generation of isogenic UAS-UBQLN2 stocks using PhiC31 integration was previously described. See Table supplement 2 for fly stocks used.

### Genetic screening

We employed the Bloomington Deficiency Kit for chromosome 2 (DK2L and DK2R) comprised of 194 different lines. All 194 lines were crossed to GMR-Gal4/Cyo or homozygous GMR>UBQLN2^P497H^ or GMR>UBQLN2^4XALS^ flies at 29°C. A minimum of 30 F1 progeny containing GMR>UBQLN2 either the Df chromosome or balancer chromosome were analyzed for eye morphology 1-3 days post eclosion using a blinded, 1-5 grading system, with a score of 1 representing a control (GMR-Gal4) eye; 3 corresponding to the unmodified UBQLN2^4XALS^ phenotype at 29°C; and 5 representing severe eye degeneration featuring more than 50% necrotic tissue. We were unable to derive UBQLN2 progeny for a handful Df lines crossed to UBQLN2^4XALS^, suggesting lethal genetic interactions. All putative modifier Dfs were retested in secondary screens that included side-by-side crosses to GMR-Gal4, GMR>UBQLN2^WT^, GMR>UBQLN2^P497H^ and GMR>UBQLN2^4XALS^. Those Dfs that were confirmed to modify GMR>UBQLN2 eye phenotypes in both screens were deemed bona fide modifier Dfs. Sexually dimorphic phenotypes were also scored.

### Drosophila climbing assay

Climbing assay was modified from methods described previously^31^. Climbing ability was measured by tapping ∼10 flies to the bottom of a graduated testing vial (15 cm) and taking videos over of fly movement over the course of 10 sec. More than 100 flies for each genotype and each gender were used for climbing ability. Video frames at the 5 second time point were used to record the position of each fly using the multi-point plugin in ImageJ. Using the final positions of every fly and respective starting points also marked with ImageJ’s multi-point, the verticle displacement and velocity of every fly was calculated. Data showing velocity of each individual fly were graphed as scattered plots with mean climbing distance and SEM. Unpaired t-test with Welch’s correction were used for statistical analysis for different groups of flies.

### NMJ assay

NMJ assay was modified from methods described previously^31^. Third-instar, wandering larvae from the F1 generation were rinsed in ice-cold PBS (Lonza, 17-512F) and dissected along the dorsal midline. All tissues except the brain and nerves were removed to expose the muscles and NMJs. The dissected larval pelt was fixed in 4% paraformaldehyde for 20 min at room temperature. The larval pelts were given a wash with PBS followed by blocking with 5% Normal Goat Serum (NGS) in 0.1% PBST (0.1% TX-100 in PBS). Following blocking, the larval pelts were probed with primary antibodies overnight at 4°C. They were then washed several times with 0.1% PBST followed by incubation with secondary antibodies for 2 h at room temperature, subsequently followed by washes with 0.1% PBST. Larvae were then mounted onto slides using Prolong Gold mounting media. Confocal images were acquired using Zeiss LSM 710 confocal microscope and a 60x oil objective was used to image the NMJs. Both primary and secondary antibody solutions were prepared in 5% NGS in 0.1% PBST. For primary antibodies, the following dilutions were used: 1:100 Cy3-conjugated goat anti-HRP (Jackson ImmunoResearch, 123-165-021); 1:100 mouse anti-DLG 4F3 (DSHB). For secondary antibodies, the following antibody dilutions were used: 1:250 Alexa Fluor 647-conjugated phalloidin (Invitrogen, A22287); 1:500 goat anti-mouse Alexa Fluor 488 (Invitrogen, A-11029). For the analyses, NMJs innervating muscle 4 on segments A2-A3 were imaged and analyzed for synaptic bouton quantification. Mature boutons are defined as boutons that are included in a chain of 2 or more boutons. Satellite boutons are defined as a single bouton that is not included in a chain of boutons, and instead, sprout off of a mature bouton or branch. The groups were compared using Unpaired Student t-test on GraphPad Prism software. P-value less than 0.05 was considered statistically significant.

### iPSC culture and motor neuron differentiation

A normal iPSC line (WC031i-5907-6, fibroblasts from neonatal male) was obtained from WiCell Research Institute^71^. Into this line we introduced the following mutations using CRISPR/CAS9: P497H, 2XALS (P497H, P525S), 4XALS (P497H, P506T, P509S, P525S), and I498X, which harbors a 1 nt deletion in codon 497 that leads to frameshift and translation termination at codon 498. Edited iPSC lines were karyotyped, sequenced for the top five ranking off-target cleavages (none were found) and confirmed for expression of pluripotency markers. iPSCs were cultured with mTeSR1 (Stemcell Technologies) on Matrigel (Corning). iPSCs on Matrigel were passaged with 0.5 mM EDTA. iPSC colonies were passaged every 4–7 days at a 1:3 to 1:6 split ratio.

Differentiation of iPSCs into motor neurons was carried out as previously described^51^. In brief, iPSCs were dissociated and placed Matrigel coated plates. On the following day, the iPSC medium was replaced with a chemically defined neural differentiation medium, including DMEM/F12, Neurobasal medium at 1:1, 0.5xN2, 0.5xB27, and 1xGlutamax (all are from Invitrogen). CHIR99021 (3 μM, Torcris), 2 μM DMH-1 (Torcris) and 2 μM SB431542 (Stemgent) were added in the medium. The culture medium was changed every other day. Human iPSCs maintained under this condition for 7 days were induced into neuroepithelial progenitors (NEP). The NEP cells were then dissociated with Dispase (1 mg/ml) and split at 1:6 with neural differentiation medium described above. Retinoic acid (RA, 0.1 μM, Stemgent) and 0.5 μM Purmorphamine (Stemgent) were added in combination with 1 μM CHIR99021, 2 μM DMH-1, and 2 μM SB431542. The medium was changed every other day. NEP cells maintained under this condition for 7 days differentiated into OLIG2+ motor neuron progenitors (MNP). To induce motor neuron differentiation, OLIG2+ MNPs were dissociated with EDTA (0.5 mM) and cultured in suspension in the above neural differentiation medium with 0.1 μM RA and 0.1 μM Purmorphamine. The medium was changed every other day. OLIG2+ MNPs under this condition for 6 days differentiated into HB9+ NMPs. The HB9+ NMPs were then dissociated with Accutase (Invitrogen) into single cells and plated on Matrigel coated plates. The HB9+NMPs were cultured with 0.1 μM RA, 0.1 μM Purmorphamine and 0.1 μM Compound E (Millipore) for 6 days to mature into CHAT+ motor neurons.

### Microscopy

Drosophila eye pictures were acquired using Leica S9 i Stereomicroscope. For immunostaining, iPSCs and iMNs were fixed with 4% paraformaldehyde in PBS, permeabilized with 0.2% PBST, blocked with 2% BSA, and stained with primary antibodies for overnight at 4°C and the stained with α-rabbit-Alex-488 and α-mouse-Alexa-594-conjugated secondary antibodies. Images were acquired using a Nikon A1 confocal microscope using either a 20x lens or a 60x oil lens. For Lysotracker assays, iPSCs were incubated with 70 nM LysoTracker™ Red DND-99 (L7528, Invitrogen) for 1 h at 37°C in culture medium. Live microscopy of lysosomes was performed using a Nikon A1 confocal microscope with a heated chamber and an objective to maintain the cells at 37°C using a 60x oil lens.

### Immunoblotting

Fly lysates were prepared by homogenizing 10 fly heads in 50 μl of RIPA lysis buffer. The soluble fraction was taken after centrifugation at 21,000 x g in a microcentrifuge for 10 min iPSC extract were prepared using in lysis buffer containing 20 mM Tris-HCl (pH 8.0), 138 mM NaCl, 10 mM KCl, 1 mM MgCl_2_, 1 mM EDTA and 1% Triton-X 100 v/v (TX buffer). All lysis buffers were supplemented with protease inhibitor cocktail (Sigma, P8340), 10 mM NaF, and 1 mM DTT. Following centrifugation at 21,000 x g for 15 min, the insoluble pellet was washed twice with PBS, then suspended and boiled in Laemmli buffer. For immunoblotting, samples were separated by SDS-PAGE and transferred to PVDF membranes and immunoblotted with primary antibodies and LI-COR IRDye secondary antibodies (IRDye 800CW goat anti-rabbit and IRDye 680RD goat anti-mouse) as described. Signals were acquired using Odyssey bio-systems (LI-COR Biosciences). Immunoblotting results were analyzed and organized with ImageStudio Lite software (LI-COR).

### Limited proteolysis

iPSCs were harvested and lysed with lysis buffer containing 20 mM Tris-HCl (pH 8.0), 138 mM NaCl, 10 mM KCl, 1 mM MgCl_2_, 1 mM EDTA and 0.2% NP-40 v/v. A total of 20 μg of each protein lysate in 10 μl was digested with increasing amounts of chymotrypsin (0.05 μg/ml to 0. 2 μg/ml, final volume, Sigma) for 5 min at room temperature. Digestion was terminated by addition of Laemmli loading dye and boiling at 95°C for 5 min. The digested proteins were analyzed by Western blotting with anti-UBQLN2 antibodies.

### Sholl analysis

Sholl analysis was done by determining the number of neurite branches at various radial distances from the cell body. The rate at which branching decreases as a function of distance is the Sholl regression coefficient, or Sholl decay. Digitizing individual neurite branching patterns, or tracing, was performed using the Simple Neurite Tracer (SNT) plugin in ImageJ. Semi-automatic tracing of β-Tubulin staining in iMNs was carried out blinded on individual neurons. SNT’s Python application programming interface was implemented to measure the number of branch points and their respective radial distances. Other morphological descriptors including neurite length, branching number and primary neurite number, were measured in a similar way and graphed as scatter plots in Graphpad. Using the Python programming language, linear regression was fitted to the semi-log plots of the total branch points for every 1 μm radial distances.

There are a variety of ways to perform curve fitting for Sholl analysis, including linear mixed models. However, mixed models are necessary for samples with extensive clustering or heterogenity^72^. Since each sample was differentiated, immunostained and imaged simultaneously and in the same way, a simple linear regression model was appropriate. To ensure each linear fit was an accurate approximation of branch point distribution, only linear fits with a high Pearson correlation (R^2^>0.8) were used to calculate the sholl coefficient (Equation 1). Individual neuron Sholl decay was graphed as scatter plots using Graphpad, where each neuron was one data point. Since neurons with a low Pearson correlation were excluded in this analysis, all neurons of a given genotype were combined into one sholl plot, in an additional, more inclusive analysis. This was accomplished by plotting the average number of branch points at 1μm intervals amongst all neurons of a given genotype. Due to high variation of branching within each sample, only branch points that fell within the 10-90 percentile were linear fitted to yield the overall sholl decay for each genotype.

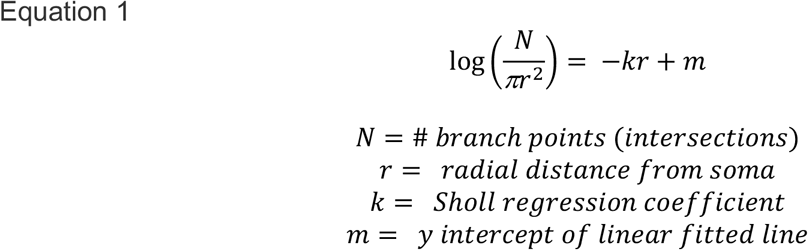

### Growth Cone Analysis

Neurite terminals to the cell body were classified as growth cones (filopodial and lamellipodial ends, Figure 9B arrows in UBQLN2^WT^ and UBQLN2^P497H^) or blunt ends (Figure 9B UBQLN2^4XALS^). The percent of neurite terminals classified as growth cones was measured for individual neurons.

### Statistical processing

Statistical analysis information including individual replicates and biological replicates number, mean or median, and error bars are explained in the Figure legends. The statistical tests and resulting *p* values are showed in the Figure legends and/or Figure panels.

## Acknowledgments

We thank Lance A Rodenkirch in optical imaging core for assistance. This work was supported by grants from the National Cancer Institute [R01CA180765-01 to R.S.T.]; National Institute of Neurological Disorders and Stroke [1R21NS090313-01A1 to R.S.T.]; National Institute on Aging [R21 AG065896-01A1 to S.H.K.]; University of Wisconsin Carbone Cancer Center Support Grant [P30CA014520]; NIH Shared Instrumentation Grants [1S10RR025483-01].

## Author contributions

S.H.K. and R.S.T. conceived the project and designed the experiments. S.H.K. and K.D.N. performed iPSC culture and differentiation experiments, molecular and biochemical assays, fixed and live-cell imaging, and Drosophila experiments. K.D.N. contributed to Sholl analysis and growth cones analysis. Y.L., N.R., and C.J.K. performed Drosophila screening, and E.N.A performed NMJ assay with support of U.B.P. W.J. analyzed RNA sequencing data, and M.A.S. performed MS experiments and analysis with support of L.M.S. S.H.K. and K.D.N. developed the figures and S.H.K. and R.S.T interpreted the data and wrote the manuscript.

## Data availability

The data that supports the findings of this study are available from the corresponding author upon request.

## Notes

### Competing Interest Statement

The authors have declared no competing interest.

