## Supplementary Figures for "Axon guidance pathways modulate neurotoxicity of ALS-associated UBQLN2"

[1]Department of Human Oncology
University of Wisconsin School of Medicine and Public Health
1111 Highland Ave.
Madison, WI 53705

[2]Department of Chemistry
University of Wisconsin-Madison
1101 University Ave.
Madison, WI 53706

[3]Department of Pediatrics, Children's Hospital of Pittsburgh
University of Pittsburgh Medical Center
Pittsburgh, PA 15224

Kim et al Figure supplement 1

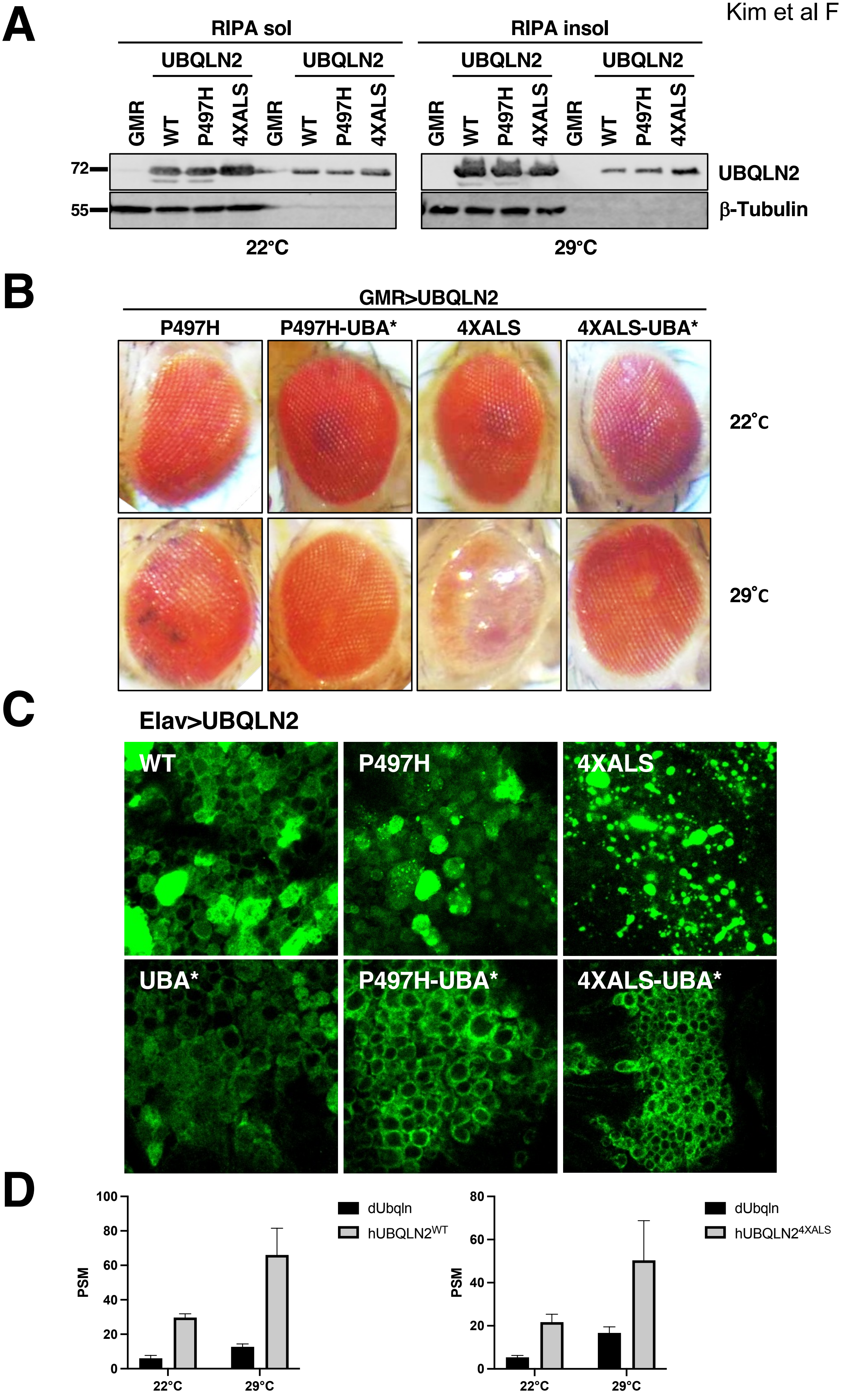

**Figure supplement 1**. **Ub-binding contributes to eye degeneration and aggregation of UBQLN2$^{ALS}$ mutants**. **(A)** Expression and RIPA solubility of UBQLN2 proteins in head extracts prepared from homozygous GMR>UBQLN2 flies of the indicated genotype. **(B)** Representative eye images from homozygous GMR>UBQLN2 flies of the indicated genotype reared at 22°C and 29°C. UBA* denotes an F594A mutation in the UBA domain that diminishes Ub binding. **(C)** Localization patterns of wild-type and ALS-mutant UBQLN2 proteins expressed under control of an Elav driver in the Drosophila brain. Note reduced aggregation of UBQLN2$^{4XALS\text{-}UBA*}$ relative to UBQLN2$^{4XALS}$. **(D)** Relative abundance of Drosophila Ubqln (dUbqln) and human UBQLN2 (hUBQLN2) in GMR>UBQLN2 flies. Head extracts from flies of the indicated genotypes were analyzed by mass spectrometry to determine peptide spectral matches (PSM) for dUbqln and hUBQLN2 at 22°C and 29°C. N=100 flies per genotype. The bars represent mean with *SEM* of triplicate.

Kim et al Figure supplement 2

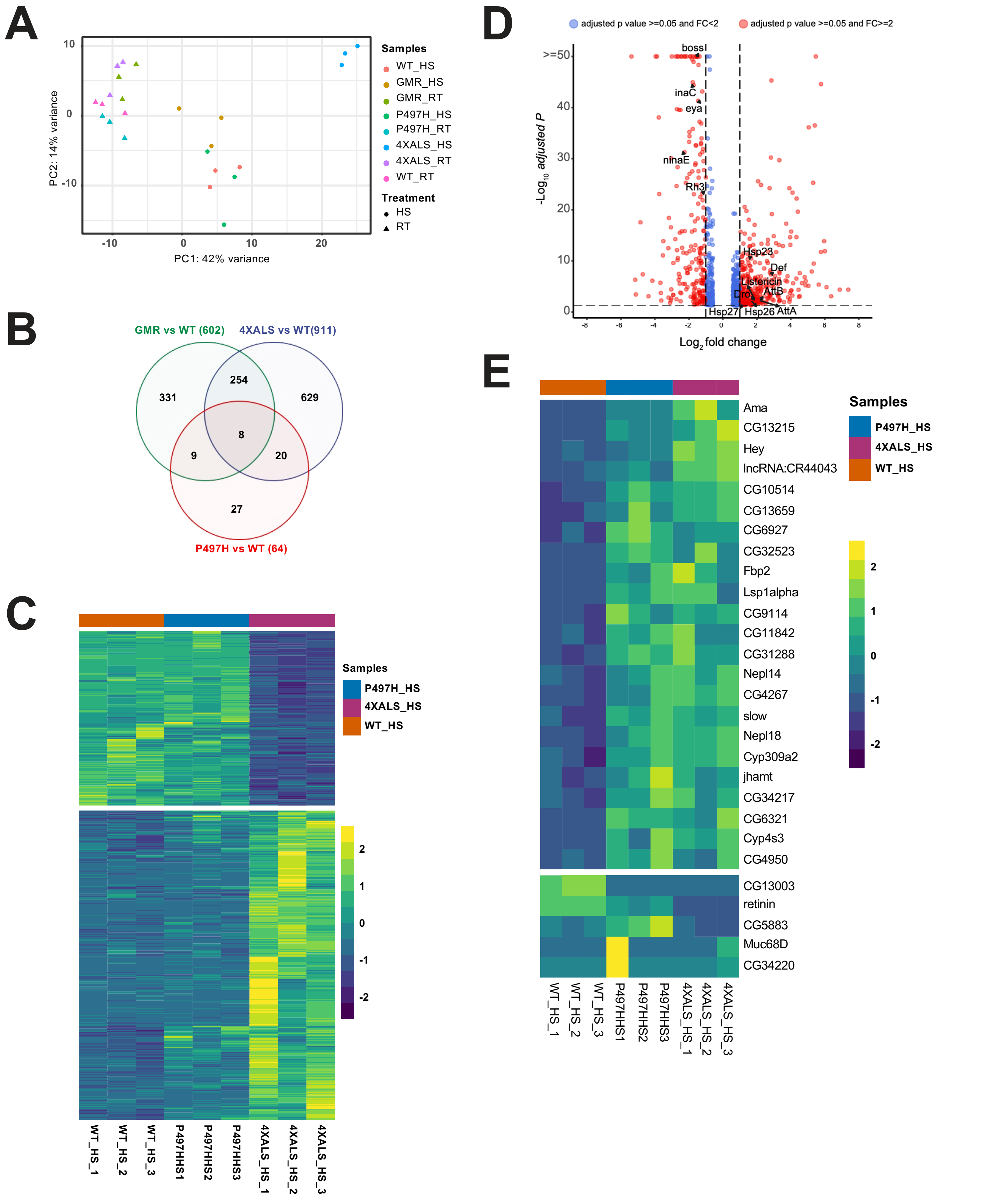

**Figure supplement 2**. **Gene expression profiles of GMR>UBQLN2$^{ALS}$ flies. (A)** Principle component analysis of RNA-Seq data from whole heads of GMR, GMR>UBQLN2$^{WT}$, GMR>UBQLN2$^{P497H}$, and GMR>UBQLN2$^{4XALS}$ flies reared at 22°C or 29°C. **(B)** Venn diagram of differentially expressed genes (DEGs) from comparisons among GMR-Gal4 control, GMR>UBQLN2$^{WT}$, GMR>UBQLN2$^{P497H}$ and GMR>UBQLN2$^{4XALS}$ flies reared at 29°C. **(C)** Heat map of genes differentially expressed between GMR>UBQLN2$^{WT}$ and GMR>UBQLN2$^{4XALS}$ fly heads at 29°C. **(D)** Volcano plot depicting genes differentially expressed between GMR>UBQLN2$^{WT}$ and GMR>UBQLN2$^{4XALS}$ flies at 29°C. Downregulated photoreceptor genes, upregulated innate immunity genes, and upregulated small HSPs are highlighted. **(E)** Differentially expressed genes common to GMR>UBQLN2$^{P497H}$ and GMR>UBQLN2$^{4XALS}$ flies in comparison to GMR>UBQLN2$^{WT}$ flies at 29°C.

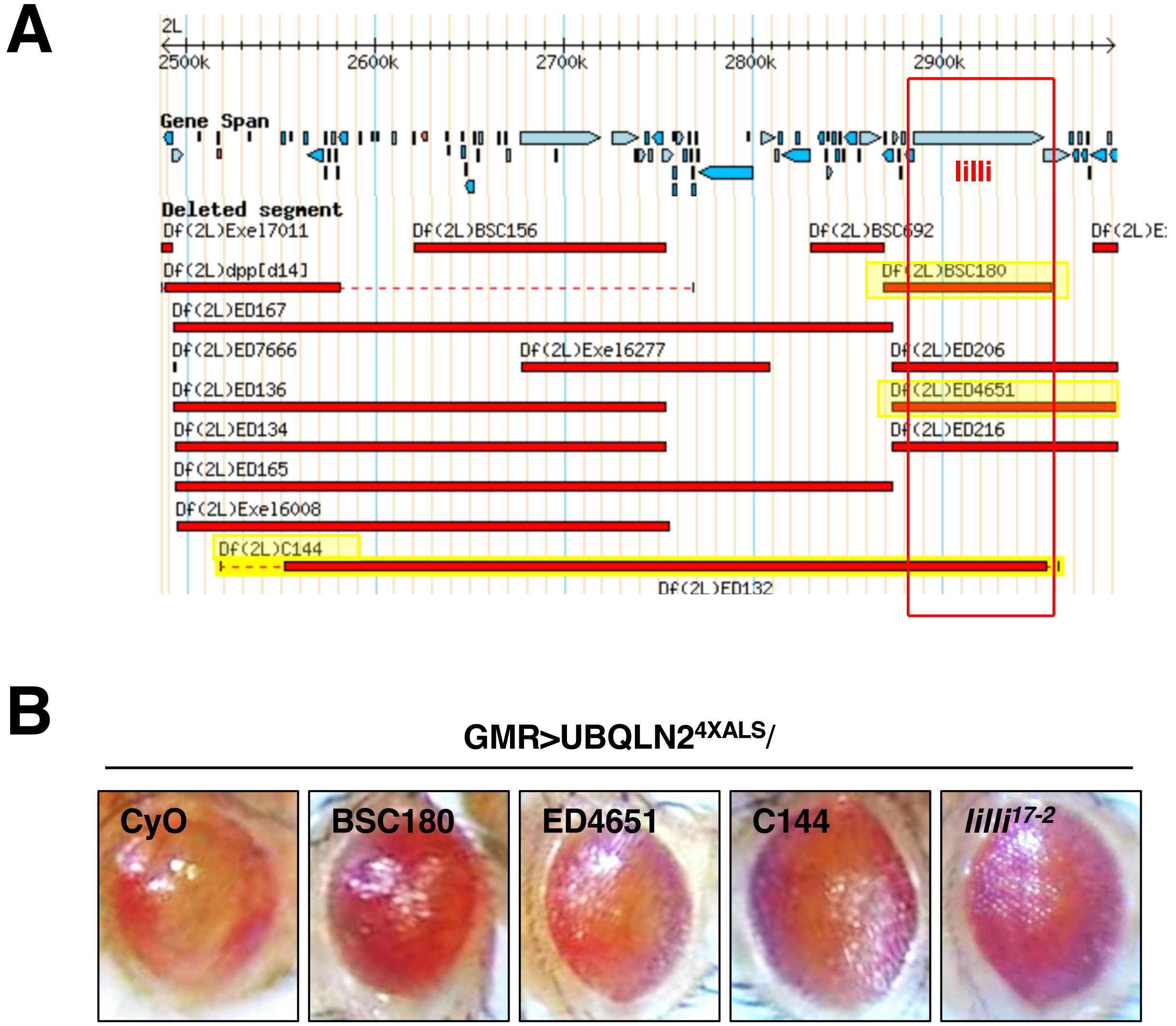

**Figure supplement 3. Transcriptional elongation factor *lilli* is a UBQLN2$^{4XALS}$ suppressor. (A)** Schematic of BSC180, ED4651, C144, and the *lilli* gene locus. **(B)** Representative eye phenotypes from GMR>UBQLN2$^{4XALS}$ flies harboring the indicated alleles.

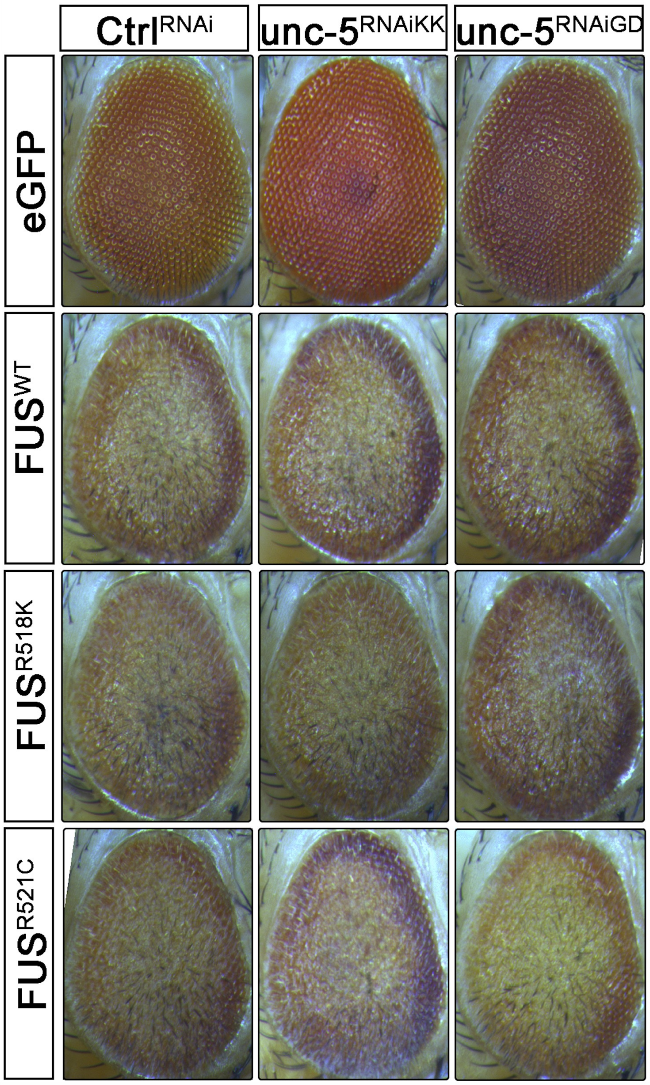

**Figure supplement 4**. ***Unc-5* silencing does not modify FUS-associated eye degeneration.** Representative eye images of GMR>eGFP (control), GMR>$FUS^{WT}$, GMR>$FUS^{R518K}$, or GMR>$FUS^{R521C}$ flies harboring the indicated shRNA alleles.

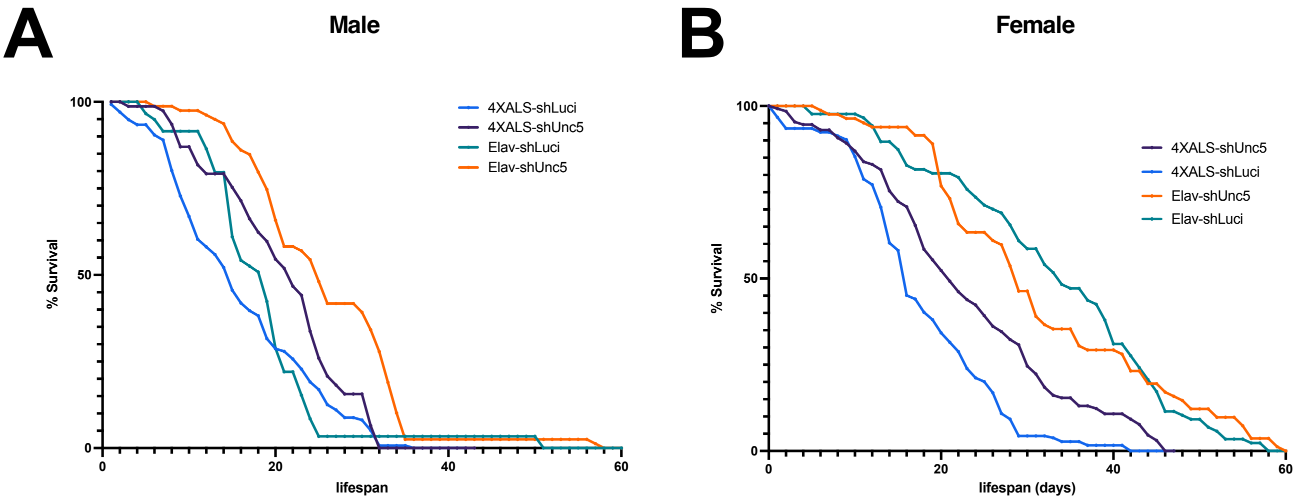

**Figure supplement 5**. **Pan-neuronal Unc-5 knockdown enhances lifespan of Elav>UBQLN2$^{4XALS}$ flies**. Recombinant Elav>UBQLN2$^{4XALS}$ or a Elav>Gal4 flies were crossed to the indicated RNAi lines (shLuci or shUnc5). Lifespan of male **(A)** and female **(B)** progeny reared at 27°C was measured as described in supplementary Methods. n>100 flies.

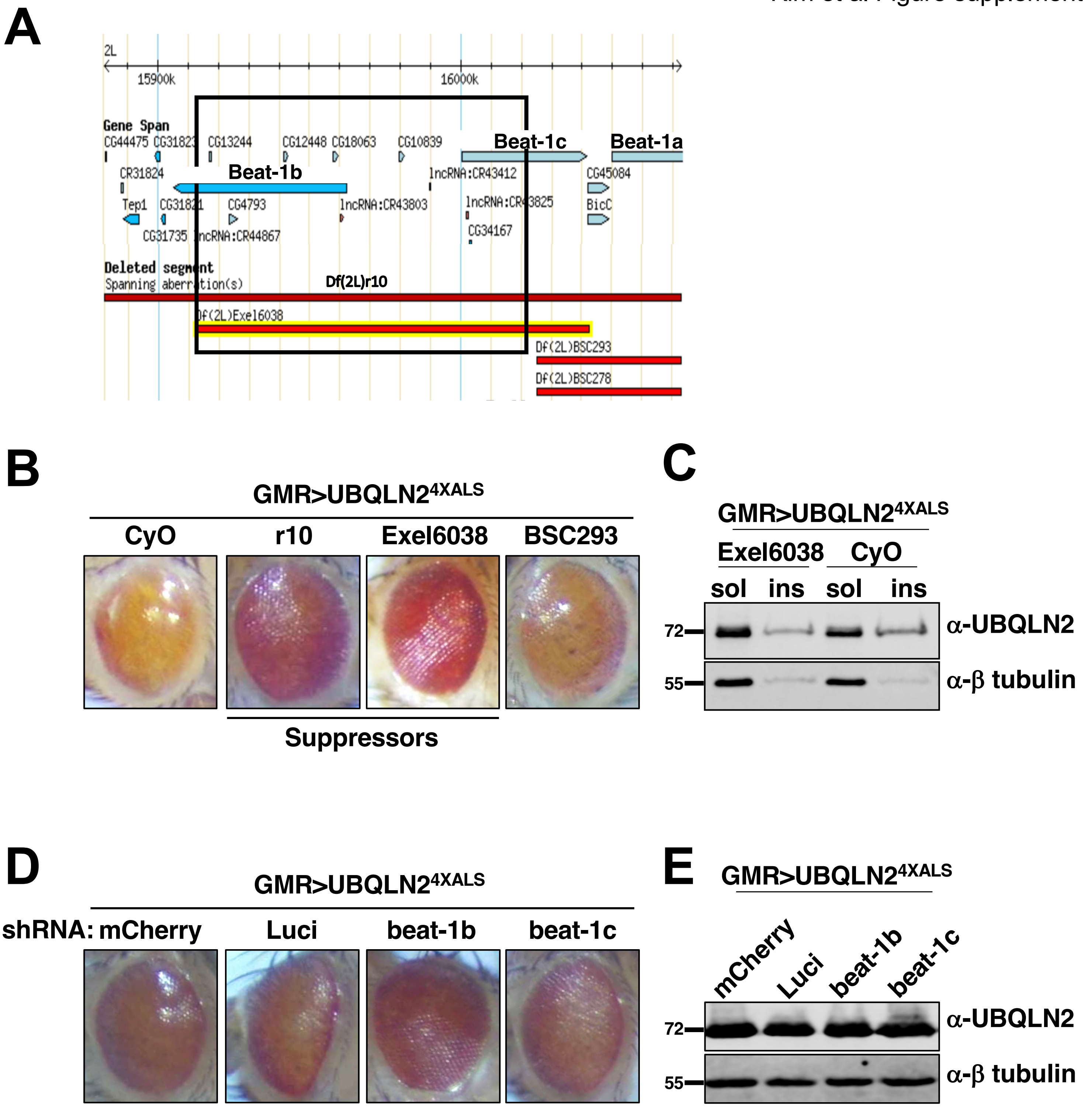

**Figure supplement 6. The axon guidance gene *beat-1b* is a UBQLN2$^{4XALS}$ suppressor.** **(A)** Schematic of the *beat-1b* gene locus. **(B)** Representative eye phenotypes of GMR>UBQLN2$^{4XALS}$ flies harboring the indicated Df alleles. **(C)** UBQLN2 expression levels and RIPA solubility in whole heads of GMR>UBQLN2$^{4XALS}$/Exel6038 and GMR>UBQLN2$^{4XALS}$/Cyo flies. **(D)** Representative eye phenotypes of GMR>UBQLN2$^{4XALS}$ flies expressing the indicated shRNAs. **(E)** Knockdown of Beat-1b or Beat-1c does not inhibit UBQLN2 expression in GMR>UBQLN2$^{4XALS}$ flies.

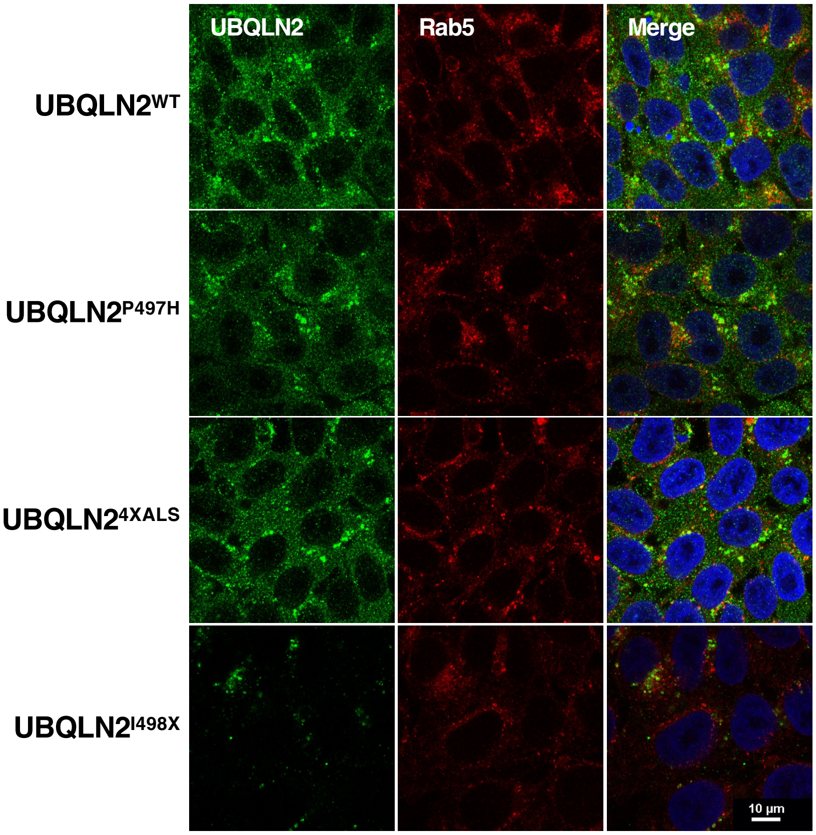

**Figure supplement 7. Endogenous UBQLN2 proteins are not colocalized with early endosomes after BafA1 treatment**. UBQLN2$^{WT}$, UBQLN2$^{P497H}$, UBQLN2$^{4XALS}$, and UBQLN2$^{I498X}$ iPSCs were immunostained with UBQLN2 and Rab5 antibodies after incubation in 100 nM BafA1 for 16 h.

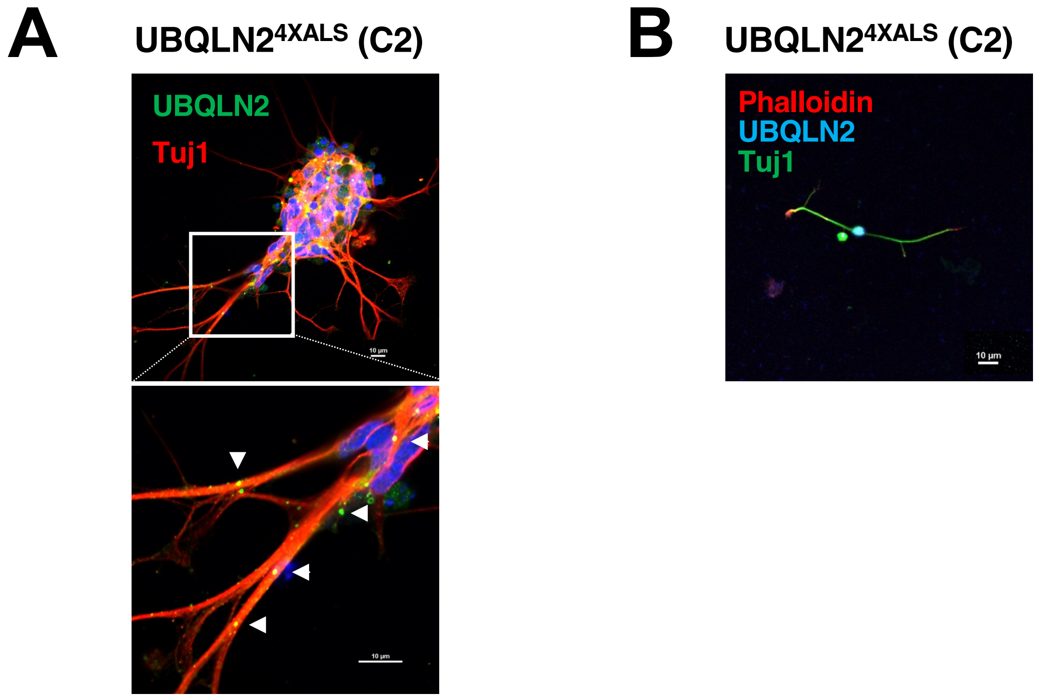

**Figure supplement 8. Protein aggregation and reduced neurite complexity in an independent UBQLN2$^{4XALS}$ iMN line. (A)** UBQLN2$^{4XALS}$ (C2) iMNs were immunostained for UBQLN2 and Tuj1. Aggregates are marked with arrowheads. **(B)** Growth cone morphologies in UBQLN2$^{4XALS}$ (C2) iMNs. iMNs were stained with α-UBQLN2, α-Tuj1, and phalloidin. Note reduced growth cone elaboration in UBQLN2$^{4XALS}$ (C2) iMNs.

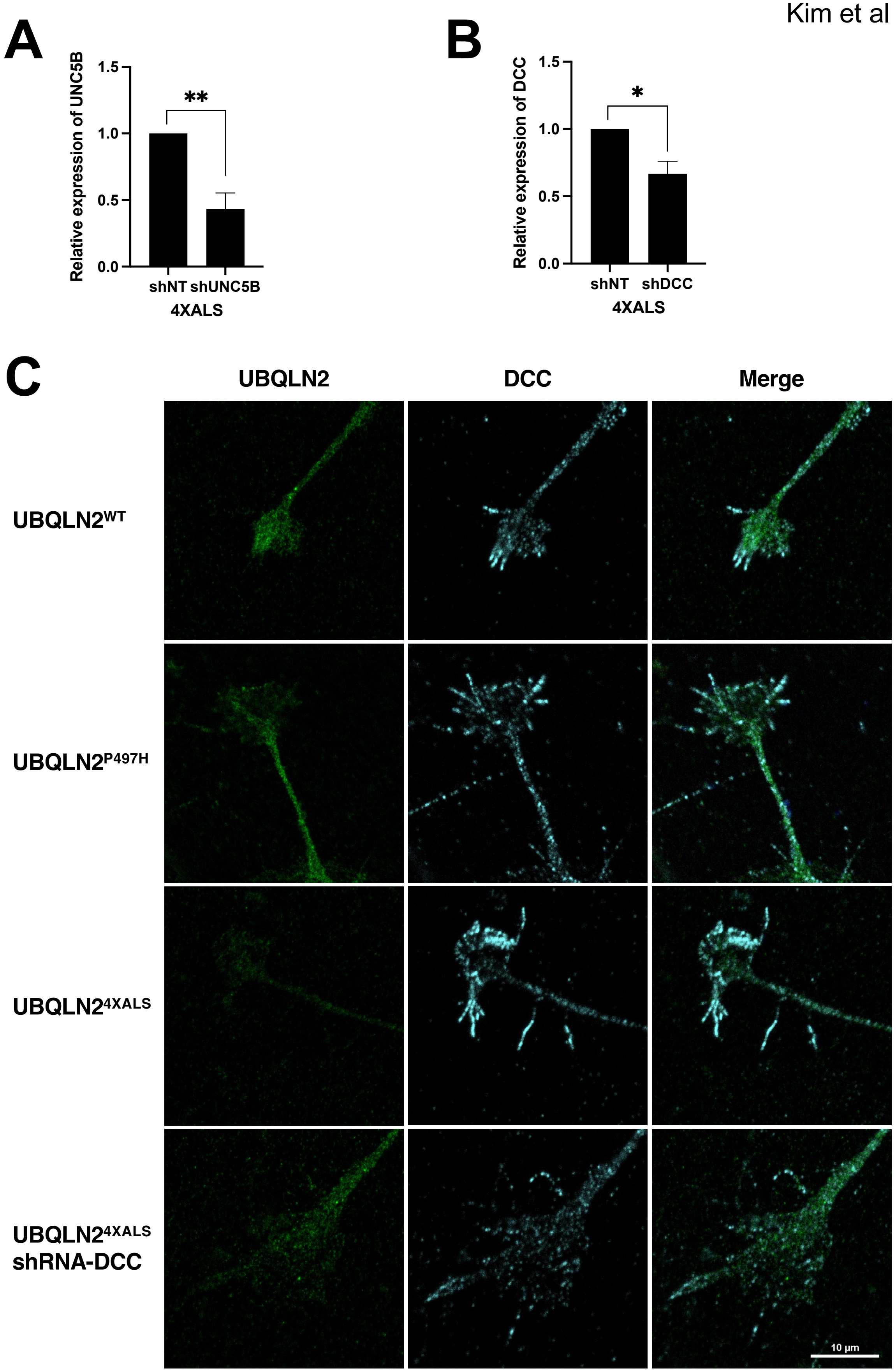

**Figure supplement 9. Expression of UNC5B and DCC mRNA in UBQLN2$^{4XALS}$ iPSCs transduced with lentiviral shRNA vectors.** mRNA levels of UNC5B **(A)** and DCC **(B)** were analyzed by RT-qPCR in iPSCs expressing the indicated shRNAs. The bars represent mean with *SEM* of triplicate. **(C)** iMNs of the indicated genotypes were immunostained with $\alpha$-DCC antibodies. Note reduced filopodial DCC signal intensity in UBQLN2$^{4XALS}$: shDCC iMNs.

**Table supplement 1. Modifiers for GMR>UBQLN2**

| BDSC # | Deficiency | Suppressor/Enhancer | |
|---|---|---|---|
| | | GMR>UBQLN2$^{4XALS}$ | GMR>UBQLN2$^{P497H}$ |
| 6609 | BSC19 | Suppressor | |
| 9610 | BSC180 | Suppressor | |
| 23677 | BSC292 | Suppressor | |
| 8904 | ED4651 | Suppressor | |
| 9064 | ED2426 | Suppressor | |
| 7521 | Exel6038 | Suppressor | |
| 90 | C144 | Suppressor | |
| 7144 | BSC37 | Enhancer | Enhancer |
| 9507 | BSC148 | Enhancer | |
| 24378 | BSC354 | Enhancer | |
| 24132 | ED629 | Enhancer | |
| 24133 | ED690 | Enhancer | |
| 24113 | ED1102 | Enhancer | |
| 9682 | ED1378 | Enhancer | |
| 24132 | ED629 | Enhancer | |
| 9270 | ED250 | No effect | Enhancer |
| 24652 | ED441 | No effect | Enhancer |
| 8910 | ED2219 | No effect | Enhancer |
| 9278 | ED2747 | No effect | Enhancer |
| 7546 | Exel6064 | Lethal enhancer | Enhancer |
| 442 | CX1 | Lethal enhancer | |
| 741 | M41A10 | Lethal enhancer | |
| 1702 | X1, Mef2[X1] | Lethal enhancer | |
| 5330 | ed1 | Lethal enhancer | |
| 6780 | 14H10W-35 | Lethal enhancer | |
| 7783 | Exel7011 | Lethal enhancer | |
| 7888 | Exel7144 | Lethal enhancer | |
| 7896 | Exel7162 | Lethal enhancer | |
| 8673 | BSC107 | Lethal enhancer | |
| 8674 | BSC109 | Lethal enhancer | |
| 9641 | BSC213 | Lethal enhancer | |
| 9716 | BSC241 | Lethal enhancer | |
| 23152 | BSC252 | Lethal enhancer | |
| 25428 | BSC595 | Lethal enhancer | |
| 26782 | lt109 | Lethal enhancer | |

**Table supplement 2. Drosophila stocks used in this study**

| Gene/nature of allele | Genotype | SOURCE | IDENTIFIER/REF |
|---|---|---|---|
| Deficiency | w[1118]; Df(2L)Exel6038, P{w[+mC]=XP-U}Exel6038/CyO | BDSC | BDSC7521 |
| Beat-1c/RNAi | y[1] sc[*] v[1] sev[21]; P{y[+t7.7] v[+t1.8]=TRiP.HMC05547}attP40 | BDSC | BDSC64528 |
| Beat-1a/RNAi | y[1] sc[*] v[1] sev[21]; P{y[+t7.7] v[+t1.8]=TRiP.HMC05811}attP40 | BDSC | BDSC64938 |
| Beat-1b/RNAi | y[1] sc[*] v[1] sev[21]; P{y[+t7.7] v[+t1.8]=TRiP.HMC04226}attP40 | BDSC | BDSC55938 |
| Rab5/RNAi | y[1] sc[*] v[1] sev[21]; P{y[+t7.7] v[+t1.8]=TRiP.HMS00147}attP2 | BDSC | BDSC34832 |
| Luciferase/RNAi | y[1] v[1]; P{y[+t7.7] v[+t1.8]=TRiP.JF01355}attP2 | BDSC | BDSC31603 |
| Rab5/RNAi | y[1] v[1]; P{y[+t7.7] v[+t1.8]=TRiP.JF03335}attP2 | BDSC | BDSC30518 |
| beat-1b/P element | w[1118]; PBac{w[+mC]=WH}beat-Ib[f04746] | BDSC | BDSC18802 |
| Rab5/overexpression | w[*]; P{w[+mC]=UAS-GFP-Rab5}3 | BDSC | BDSC43336 |
| Unc-5/RNAi | y[1] sc[*] v[1] sev[21]; P{y[+t7.7] v[+t1.8]=TRiP.HMS01099}attP2 | BDSC | BDSC33756 |
| Unc-5/RNAi | NA | VDRC/GD | VDRC8138 |
| Unc-5/RNAi | NA | VDRC/KK | VDRC110155 |
| GFP/overexpression | w[1118]; P{w[+mC]=UAS-EGFP}34/TM3, Sb[1] | BDSC | BDSC5430 |
| Lilli/loss of function | lilli[A17-2] cn[1] bw[1]/CyO | BDSC | BDSC5726 |
| Deficiency | w[1118]; Df(2R)BSC346/CyO | BDSC | BDSC24370 |
| mCherry/RNAi | y[1] sc[*] v[1] sev[21]; P{y[+t7.7] v[+t1.8]=UAS-mCherry.VALIUM10}attP2 | BDSC | BDSC35787 |
| Frazzled/overexpression | y[1] w[*]; P{w[+mC]=UAS-fra}3 | BDSC | BDSC8814 |
| NetB/RNAi | y[1] v[1]; P{y[+t7.7] v[+t1.8]=TRiP.JF01882}attP2 | BDSC | BDSC25861 |
| NetA/RNAi | y[1] v[1]; P{y[+t7.7] v[+t1.8]=TRiP.JF01232}attP2 | BDSC | BDSC31288 |
| Frazzled/RNAi | y[1] v[1]; P{y[+t7.7] v[+t1.8]=TRiP.JF01231}attP2 | BDSC | BDSC31469 |
| Frazzled/RNAi | y[1] v[1]; P{y[+t7.7] v[+t1.8]=TRiP.JF01457}attP2 | BDSC | BDSC31664 |
| Frazzled/RNAi | y[1] sc[*] v[1] sev[21]; P{y[+t7.7] v[+t1.8]=TRiP.HMS01147}attP2 | BDSC | BDSC40826 |
| NetA/deletion | w[1118] NetA[Delta] | BDSC | BDSC66878 |
| NetB/deletion | y[*] w[*] NetB[Delta] | BDSC | BDSC66879 |
| Deficiency (lilli) | BSC180 | BDSC | BDSC9610 |
| Deficiency (Rab5) | BSC37 | BDSC | BDSC7144 |
| Deficiency (lilli) | ED4651 | BDSC | BDSC94697 |
| Deficiency (lilli) | C144 | BDSC | BDSC99 |
| Lilli (LOF allele) | lilli[A17-2] cn[1] bw[1]/CyO | BDSC | BDSC5726 |
| Unc-5 (LOF) | $Unc\text{-}5^{3}$ | Gregory Bashaw | [1] |
| Unc-5 (LOF) | $Unc\text{-}5^{8}$ | Gregory Bashaw | [1] |
| Unc-5/overexpression | HA-Unc-5 (Chr2) | Gregory Bashaw | [2] |
| Unc-5/overexpression | HA-Unc-5 (Chr3) | Gregory Bashaw | [2] |

### Supplementary Methods

#### Fly brain immunohistochemistry

The protocol for fly brain immunohistochemistry was adapted from previously published protocol[3]. Adult fly brains were dissected using a pair of fine forceps in PBS (or 0.3% TX-100 in PBS), fixed, blocked with normal goat serum, and stained with primary antibodies at 1:500 dilution for two overnights at 4°C. After subsequent secondary antibody staining, DAPI was added for the nuclear staining. Images were acquired using a Nikon A1 confocal microscope using a 60x oil lens.

#### Mass spectrometry

Fly lysates were prepared by homogenizing 100 male fly heads in 200 µl of lysis buffer containing 20 mM Tris-HCl (pH 8.0), 138 mM NaCl, 10 mM KCl, 1 mM $MgCl_2$, 1 mM EDTA, 0.5% sodium deoxycholate w/v, and 0.1% SDS w/v (RIPA buffer w/o NP-40). Samples were centrifuged at 20,000 x g for 10 min and soluble fractions subjected to tryptic digestion and orbitrap mass spectrometry (MS) using the filter aided sample preparation (FASP) method[4]. We performed two technical replicates for each of the three biological replicates. The tryptic digest solution was desalted/concentrated using an Omix 100 µl (80 µg capacity) C18 tip and the peptides were analyzed by HPLC-ESI-MS/MS using a system consisting of a high performance liquid chromatograph (nanoAcquity, Waters) connected to an electrospray ionization (ESI) Orbitrap mass spectrometer (QE HF, ThermoFisher Scientific). HPLC separation employed a 100 x 365 µm fused silica capillary micro-column packed with 20 cm of 1.7 µm-diameter, 130 Angstrom pore size, C18 beads (Waters BEH), with an emitter tip pulled to approximately 1 µm using a laser puller (Sutter instruments). Peptides were loaded on-column at a flow-rate of 400 nl/min for 30 min and then eluted over 120 min at a flow-rate of 300 nl/min with a gradient of 5% to 35% acetonitrile, in 0.2% formic acid. Full-mass profile scans were performed in the FT orbitrap between 375-1500 m/z at a resolution of 120,000, followed by MS/MS HCD scans of the ten highest intensity parent ions at 30% relative collision energy and 15,000 resolution, with a mass range starting at 100 m/z. Dynamic exclusion was enabled with a repeat count of one over a duration of 30 sec. The MetaMorpheus software program was used to identify peptides and proteins in the samples[5,6]. Protein fold changes were quantified by FlashLFQ[7-9].

#### RNA-Seq and gene expression

Total RNA was isolated from 100 male fly heads using the TRIzol reagent (Invitrogen, 15596018) following the manufacturer's protocol and treated with TURBO Dnase (Invitrogen, AM2239). RNA samples were prepared three biological replicates for each genotype and each temperature. Then RNA samples were sent to Novogene (Novogene Co., Ltd, Sacramento, CA) for non-stranded cDNA library building and sequencing at PE150 with NovoSeq 6000. Raw reads adapters were trimmed by fastp[10] and then were mapped to Drosophila melanogaster genome(dmel-all-chromosome-r6.27, FlyBase) by STAR with the setting suggested by ENCODE project (https://github.com/ENCODE-DCC/rna-seq-pipeline). The number of RNA-Seq reads mapped to each transcript were summarized with featureCounts[11] and differential expression was called using DESeq2[12]. The Gene ontology analysis was performed on MetaScape website[13].

#### Drosophila longevity assays

Longevity assay was modified from methods described previously[3]. For survival analysis, flies were aged at 27°C with no more than 15 flies per vial. Total more than 100 flies were used for each genotype. Vials were changed on a 2–3 day cycle. Death events were scored on a daily basis. Rescue in longevity was defined as greater than 5% increase in median lifespan in addition to the statistical threshold according to the Log-rank (Mantel-Cox) test, *p*<0.05. In the survival graphs shown, each set of experiments was done in the same time period with the corresponding control subjects in order to control longevity variation caused by environmental factors. Both genders were used in the survival assay unless otherwise specified.

**iPSC lines used in this study**

| iPSC lines | UBQN2 genotype | SOURCE |
|---|---|---|
| UBQLN2$^{WT}$ | WT | WC031i-5907-6 |
| UBQLN2$^{P497H}$ | P497H | mutation using CRISPR |
| UBQLN2$^{2XALS}$ | P497H, P525S | mutation using CRISPR |
| UBQLN2$^{4XALS}$ | P497H, P506T, P509S, P525S | mutation using CRISPR |
| UBQLN2$^{I498X}$ | I498X | mutation using CRISPR |

**Key resources table**

| REAGENT or RESOURCE | SOURCE | IDENTIFIER |
|---|---|---|
| **Antibodies** | | |
| Mouse monoclonal anti-UBQLN2 (6H9) | Abcam | Ab190283 |
| Rabbit polyclonal anti-UBQLN2 (D7R2Z) | Cell Signaling Technology | 85509 |
| Mouse monoclonal anti-β-Tubulin | Millipore | 05-661 |
| Rabbit polyclonal anti-LC3A/B | Cell Signaling Technology | 12741S |
| Mouse monoclonal anti- LAMP1 (H4A3) | Santa Cruz Biotechnology | sc-20011 |
| Rabbit polyclonal anti-Rab5A | Cell Signaling Technology | 46449S |
| Rabbit polyclonal anti-UBQLN1 | Cell Signaling Technology | 14526 |
| Mouse monoclonal anti-Tuj1 | EMD Millipore | MAB1637MI |
| Rabbit polyclonal anti-Unc5B (D9M7Z) | Cell Signaling Technology | 13851S |
| Rabbit polyclonal anti-DCC | Invitrogen | PA5-50946 |
| Alexa Fluor™ 594 Phalloidin | Invitrogen | A12381 |
| Cy3-conjugated goat anti-HRP | Jackson ImmunoResearch | 123-165-021 |
| Mouse anti-DLG | DSHB | 4F3 |
| **Chemicals** | | |
| CHIR99021 | Tocris | 4423 |
| DMH-1 | Tocris | 4126 |
| SB431542 | Stemgent | 04-0010 |
| Retinoic acid | Stemgent | 04-0021 |
| Purmorphamine | Stemgent | 04-0009 |
| Compound E | Millipore | 565790 |
| **shRNAs** | | |
| shUNC5B-1 | Sigma | TRCN0000442978 |
| shUNC5B-2 | Sigma | TRCN0000438199 |
| shDCC-1 | Sigma | TRCN0000010318 |

| | | |
|---|---|---|
| shDCC-2 | Sigma | TRCN0000039814 |
| **Primers** | | |
| GAPDH-F | GTCTCCTCTGACTTCAACAGCG | q-PCR |
| GAPDH-R | ACCACCCTGTTGCTGTAGCCAA | q-PCR |
| UNC5B-F | ACTGCCGTGACTTCGACAC | q-PCR |
| UNC5B-R | GCCTTGCCGTCTTAAAGTTGA | q-PCR |
| DCC-F | GACTTTACCAATGTGAGGCATCT | q-PCR |
| DCC-R | GGTCCTGCTACTGCAACTTTT | q-PCR |
